# Retroviral CA subdomains encode their immature assembly properties independently of strict subdomain co-pairing

**DOI:** 10.64898/2026.10.16.759032

**Authors:** M Percipalle, A Thader, VV Hodirnau, RA Dick, FKM Schur

## Abstract

Retroviral capsid (CA) proteins display a conserved two-domain architecture consisting of independently folded N-terminal (CA-NTD) and a C-terminal (CA-CTD) subdomains. Despite a strong CA protein fold conservation, different retrovirus species are employing remarkably variable geometrical solutions to stabilize the CA lattices that form the immature virus particle. For instance, while HTLV-1 has been reported to use the CA-NTD to stabilize its immature lattice, HIV-1 relies on a contiguous structural unit formed of the CA-CTD together with the short spacer peptide 1 (SP1). This raises the question whether immature assembly phenotypes are modular features encoded independently into individual CA subdomains, or if they are emergent features requiring strict co-pairing or co-evolution between CA subdomains of a given species. To test for these two options, we have designed chimeric proteins pairing the CA subdomains of HTLV-1 and HIV-1. Our results show that the CA subdomains of HIV-1 and HTLV-1 can be clearly distinguished into assembly-passive and assembly-driving categories. Moreover, pairing assembly-driving subdomains from the two retrovirus species still allows them to independently form their native immature contacts, while at the same time trying to impose their preferred curvature on each other. Mutations of either subdomain within the chimeric context do not disrupt assembly, but rather shift the phenotype balance while relieving curvature strains, demonstrating that all immature assembly properties can be encoded into a single subdomain. Our results therefore highlight that the two-domain architecture of retroviral capsid is a result of a division of labor approach built on structural modularity, where, at least for immature assembly, one of the subdomains is the key driver for assembly competence while the other could serve more regulatory functions.

## Introduction

Retroviral assembly is mediated by the Gag polyprotein, which contains the matrix (MA), capsid (CA), and nucleocapsid (NC) domains, which are responsible for membrane targeting, protein-protein interactions and viral genome binding, respectively. In some retroviruses, additional non-canonical regions exist, such as spacer peptide 1 (SP1) in Human Immunodeficiency virus 1 (HIV-1)^1^, which play important roles in regulating assembly and virus maturation. During maturation, proteolytic processing of Gag into its individual domains, induces extensive lattice rearrangements that transform the immature spherical assembly into the mature capsid core adopting a fullerene geometry^2–4^.

Although retroviruses diverge substantially in CA sequence and Gag maturation strategy, all structurally characterized retrovirus species converge onto geometrically similar lattices to form the mature core, consisting of CA hexamers and pentamers. Despite these similar mature intra- and inter-hexameric/pentameric interfaces, this results in cores of different shapes formed across different retrovirus species. These can adopt varying architectures, including conical, tubular, spherical or irregular shapes^3,4^.

Intriguingly, the immature hexameric CA lattices (i.e., within the context of full-length Gag) found in different retrovirus species, instead form architecturally similar spherical assemblies. However, the geometric interactions stabilizing the immature CA lattice display drastic structural variations on the quaternary protein-protein interaction level^5–10^.

This can be partially explained by the fact that although all retroviral CA proteins share a conserved two-domain architecture consisting of an N-terminal subdomain (CA-NTD) and a C-terminal subdomain (CA-CTD), different retroviruses appear to distribute their immature lattice-stabilizing interactions differently between these subdomains. In the lentivirus HIV-1, stabilization of the immature lattice has largely been attributed to the CA-CTD together with SP1 which form one contiguous structural unit. Similarly, extensive interactions in the CA-CTD as well as adjacent spacer peptides or comparable structured regions that are important for immature lattice assembly have been identified in immature Rous-Sarcoma virus (RSV), Murine Leukemia virus (MLV), Equine Infectious Anemia virus (EIAV) and Mason-Pfizer Monkey virus (M-PMV)^6,8,11–15^. Here, the CA-NTD is instead thought to play a more important role during mature capsid formation.

In contrast, recent structural studies of the deltaretrovirus Human T-cell Leukemia virus 1 (HTLV-1) have revealed an unusual immature lattice organization in which HTLV-1 CA is able to adopt a remarkably flat lattice architecture where a variable spacing between the CA lattice and membrane is visible^9,10,16^. More surprisingly, structural studies revealed that it is not the CA-CTD that stabilizes the immature lattice, but the CA-NTD via the formation of extensive intra- and inter-hexamer interactions^9,10^.

In line with the unusual assembly phenotype in HTLV-1, recombinantly purified wild type (wt) HTLV-1 CA exclusively forms tubular capsid-like particle (CLP) assemblies *in vitro* in our experiments (Fig. S1A,B) as previously reported^9^, where curvature is only achieved perpendicular to the tubular axis. A planar assembly phenotype has also been observed where HTLV-1 CA forms entirely flat sheets *in vitro*^17^. In comparison, HIV-1 CA-SP1 exclusively forms spherical CLP assemblies *in vitro* (Fig. S1C,D). This suggests that different curvature preferences are imprinted into the CA domains, with substantial functional implications for virus assembly.

Overall, these observations suggest that HIV-1 and HTLV-1 represent fundamentally distinct geometrical solutions to the same biological problem of immature retroviral lattice assembly. This is intriguing from an evolutionary point of view, since the two-domain retroviral capsid exists as a result of a gene duplication event^18^, where over time the CA-CTD retained highly conserved patches in the major homology region, and the CA-NTD diverged more strongly. Therefore, HTLV-1 and HIV-1 represent different lineages that have evolved to use the same structural scaffold in different ways. This raises a fundamental question regarding retroviral capsid evolution: are CA assembly phenotypes in immature virus particles emergent properties of intact co-evolved CA subdomains, or are these phenotypes independently encoded within the individual N- and C-terminal CA subdomains and adjacent non-canonical peptides? If assembly behavior is strongly imprinted within specific subdomains, then divergent retroviral domains may retain the ability to assemble and to impose their native geometric preferences even when combined with heterologous partners. Alternatively, if assembly requires strict co-evolution between CA-NTD and CA-CTD, then chimeric combinations should fail to assemble because of incompatibilities across retrovirus species.

Previous experiments using mammalian cell-derived full-length Gag virus-like particles (VLPs), expressing HIV-1 and HTLV-1 Gag chimeras already suggested a partial assembly competence of HIV-1 CA and HTLV-1 CA domains and subdomains in a heterologous context^19^. Similarly, chimeras generated from domain-switched Bovine Leukemia virus (another member of the deltaretrovirus genus) and HIV-1 Gag polyprotein could form VLPs in insect cells^20^. However, given the larger Gag context, it was not possible to disentangle the contributions of the MA, NC and, in HIV-1 Gag, non-canonical domains to the observed phenotypes. Neither was the molecular nature of the formed Gag lattices determined, which would have revealed if the chimeric assemblies indeed formed defined lattice interactions.

Here, we engineered chimeric CA constructs between HIV-1 and HTLV-1 for recombinant expression and purification. We then examined their CLP assembly behavior and lattice organization using *in vitro* assembly assays, cryo-electron microscopy (cryo-EM) structure determination and mutagenesis experiments where we increased the domain distance or disrupted inter- and intra-hexameric interactions in either domain. Our results show that the assembly competent HIV-1 and HTLV-1 CA subdomains are modular entities with competing geometric preferences and do not require strict co-evolution together with their respective non-assembly competent partner subdomain to achieve their immature lattice geometry.

## Results

### Chimeric CA constructs can assemble into capsid-like particles in vitro

We first designed CA chimeras that contained the CA-NTD and CA-CTD domains of HTLV-1 and HIV-1 in alternating order. In case of HIV-1 CA-CTD we also added the SP1 peptide, given its importance in stabilizing the immature lattice and in being an integral structural part at the base of the immature lattice, where it forms a 6-helix bundle together with the last residues of the CA-CTD^21–24^. We refer to these chimeras from here on as CH1 (HTLV-1 CA-NTD/HIV-1 CA-CTD-SP1) and CH2 (HIV-1 CA-NTD/HTLV-1 CA-CTD) (Fig. 1A,B). By assessing their assembly competence and CLP morphology in known immature assembly conditions for both HIV-1 CA-SP1 and HTLV-1 CA (for used conditions, see Table S1), we aimed at attributing immature assembly properties to each retroviral subdomain. We expected that CH1 would be more likely to show assembly competence as it contains the two assembly driving modules of each virus. Instead, for CH2, the outcome was unclear given that it contained the two supposedly assembly-passive modules.

**Figure 1:**
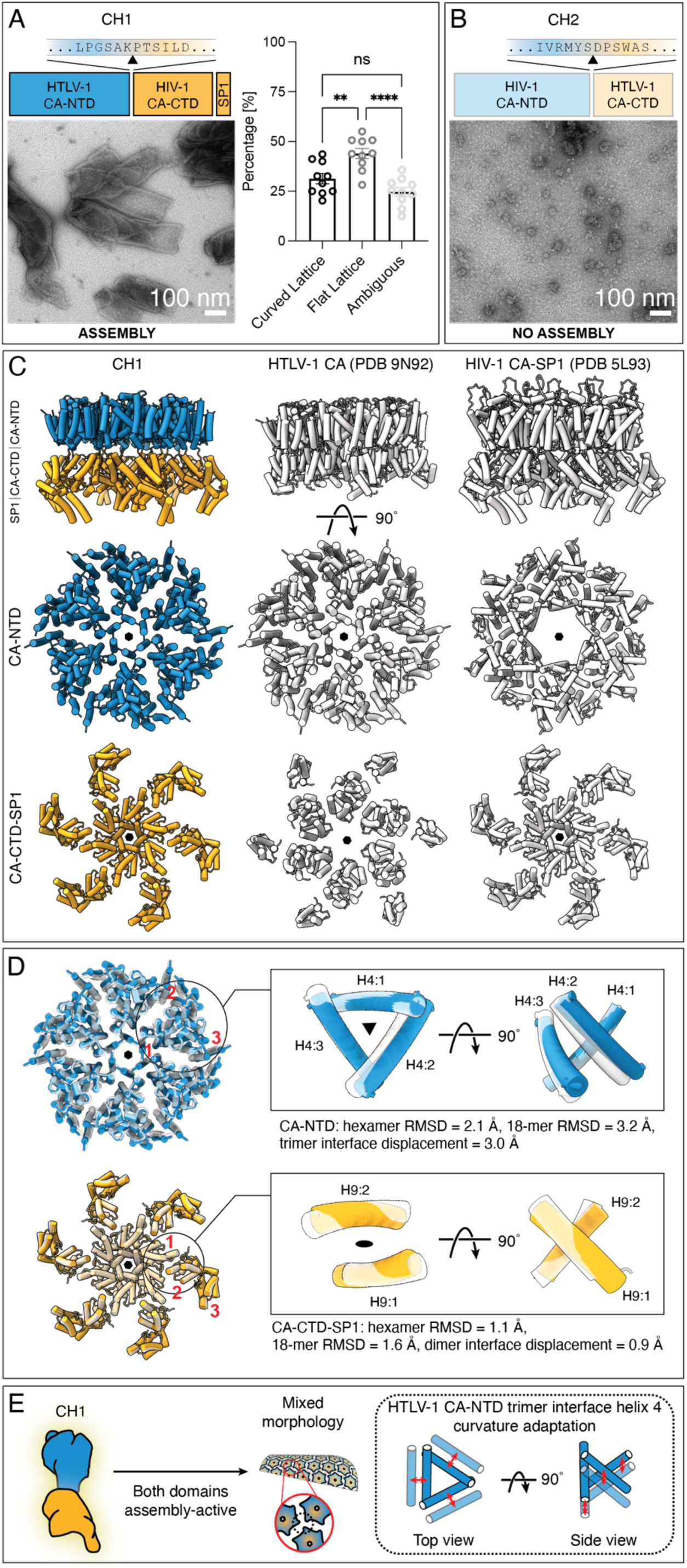
Assembly and lattice structure of HTLV-1/HIV-1 CA chimera CH1. **A)** Top: Schematic of the CH1 construct. The residues constituting the linker region connecting the HTLV-1 CA-NTD (blue) and HIV-1 CA-CTD (orange) domain are shown. Bottom: representative negative stain TEM micrograph for CH1 CLP assemblies. Right: quantification of CLP morphology, ** p = 0.0030, **** p < 0.0001. **B)** Top: Schematic of the CH2 construct, shown as in (A). Bottom: representative negative stain TEM micrograph. No CLPs are visible, only amorphous aggregates. **C)** Side-by-side comparison of the modeled CH1 Cryo-EM SPA lattice to HTLV-1 CA-NTD (PDB 9N92) and HIV-1 CA-SP1 (PDB 5L93) structures, with the CH1 CA-NTD shown in blue, the CH1 CA-CTD-SP1 shown in orange, and the wt structures in white/transparent. CH1 displays an inter-hexamer distance of 75 Å, a lattice height of 88 Å, and an inter-domain distance of 12 Å with no inter-domain interactions. While HTLV-1 CA and HIV-1 CA-SP1 have similar inter-hexamer spacings of 75 Å, HTLV-1 CA has a lattice height of 62 Å and HIV-1 CA-SP1 has a lattice height of 84 Å. Cα–Cα distances were measured in ChimeraX between adjacent hexamers to assess inter-hexamer spacing, lattice height was measured as the perpendicular distance from a least-squares plane fit through the top-layer Cα atoms to the centroid of the bottom-layer Cα atoms, and interdomain distance was measured as the Cα–Cα distance between the closest residues being part of a helix in each domain (A128 in helix 6 of the HTLV-1 CA-NTD and S175 in helix 8 of the HIV-1 CA-CTD). **D)** Alignment of an 18-mer of CH1 with the respective 18-mers of HTLV-1 CA-NTD (RMSD 3.2 Å) and HIV-1 CA-CTD-SP1 (RMSD 1.6 Å). The same RMSD measurements for the central hexamer are reported for comparison (RMSDs 2.1 and 1.1 Å, respectively). The inter-hexameric interfaces display displacements of 3.0 Å in the CA-NTD trimer interface and 0.9 Å in the CA-CTD dimer interface, highlighting how the overall lattice diverges most in the CA-NTD trimer interface. The coloring is identical to (C). **E)** Schematic explanation of how the chimeric CLP assembly arises. When both CA subdomains are assembly-active, they both impose their native contacts and preferred curvature, leading to a mixed morphology assembly. In this arrangement, the HTLV-1 CA-NTD adapts to curvature by dilation of its trimer interface. All residue numbering in CH1 is using the respective HTLV-1 CA-NTD and HIV-1 CA-CTD-SP1 numbers as shown in Figure S1A and S1B, respectively.

Indeed, CH1 formed particles under HIV-1 assembly conditions^25^ (Table S1) as assessed by negative stain TEM analysis (nsTEM) (Fig. 1A). These particles were often incomplete, adopting a bullet-like shape, and displaying a mixture of slightly more abundant tubular and flat lattice regions in their center (44% of quantified lattice) and less abundant, curved lattice regions at their tips (31%). However, this mixed morphology also resulted in a rather uncontrolled assembly with CLPs layering onto each other forming clusters where the lattice type could not unambiguously be determined (25%). The incompleteness of most CLPs appeared to stem from the flat/tubular morphology suppressing successful lattice closure at the CLP ends. This observation suggests a competitive assembly of flat vs. curved preference, where the tubular assembly, presumably imposed by the HTLV-1 CA-NTD subdomain, is exerting a stronger effect.

Under HTLV-1 immature assembly conditions^9^ (Table S1), CH1 did not assemble as efficiently and hence was not subjected to further analysis. For CH2, none of the tested assembly conditions resulted in regularly formed CLPs or other ordered structures, as only amorphous aggregates could be observed (Fig. 1B). This confirms that both the HIV-1 CA-NTD and HTLV-1 CA-CTD remain assembly-passive in this immature assembly setting.

Hence, our engineered CA chimeras show that retroviral CA subdomains can assemble into CLPs even outside of their native CA context. Furthermore, the chimeras show that there are indeed assembly-active and assembly-passive domains, and that retroviral CA subdomains do not require a co-evolved partner to perform an assembly function. The tested assembly-active domains have intrinsically strong assembling capacity capable of imposing their preferred geometry in a heterologous system, where the HTLV-1 CA-NTD is slightly more successful over HIV-1 CA-CTD-SP1 based on lattice curvature preference.

### SPA Cryo-EM structure of CH1 reveals domain independence

Next, we aimed to determine which molecular lattice interactions were formed within the CH1 CLPs and if the two assembly competent subdomains actually adopted their respective wt interactions or a completely new type of assembly. To this end, we performed single-particle analysis (SPA) cryo-EM of vitrified CH1 CLPs and determined a structure of the CH1 lattice (Fig. S2, Table S2). The global resolution of the obtained map was estimated at ∼4.1 Å FSC and while not fully resolving the helical pitch or side-chain densities, except some large aromatic residues, the map still allowed unambiguous modeling of the respective CA-NTD and CA-CTD-SP1 domains into the EM-density with high precision. The obtained structure revealed a hexameric lattice, with a hexamer-hexamer spacing of 75 Å, in line with previously measured dimensions of immature CA lattices^26,27^. The lattice had a height of 88 Å where the CA-NTD and CA-CTD were more distantly spaced compared to the wt HIV-1 CA-SP1 and HTLV-1 CA lattices (Fig. 1C).

The upper layer accommodated the HTLV-1 CA-NTD which formed lattice interactions that were essentially identical to the ones previously reported for the immature HTLV-1 CA-NTD lattice in Gag VLPs^9,10^. Intra-hexamer interactions deviated to the published wt HTLV-1 CA-NTD (PDB 9N92) by an RMSD value of 2.1 Å (Fig. 1D), while inter-hexamer interactions, measured by comparing an 18-mer CA-NTD assembly differed by 3.2 Å. Importantly, most of this deviation was due to a more plastic trimer inter-hexamer interface which had a helix displacement of 3.0 Å, which we previously identified as the key interface linking hexamers in the lattice^9^ (Fig. 1D). The displacement was measured as the average Cα shift of all remaining helices 4 after aligning one shared reference helix between wt and CH1 at each trimer interface. The lower part of the electron microscopy density could be unambiguously assigned to the immature HIV-1 CA-CTD-SP1 hexamer assembly. Intra-hexamer and inter-hexamer interactions formed by the CA-CTD and the characteristic SP1 6-helix bundle deviated to the reported wt hexamer and 18-mer (PDB 5L93) by an RMSD of 1.1 and 1.6 Å (Fig. 1D). This shows that the CA-CTD is able to assemble an immature-like arrangement independent of the HIV-1 CA-NTD arrangement *in vitro* as previously suggested^28^, and it indicates that the strong HIV-1 CA-CTD-SP1 interactions in the immature lattice do not substantially change in order to form assemblies in the heterologous assembly context. This reluctance of the CA-CTD-SP1 to change interactions to accommodate to the flat HTLV-1 CA-NTD is probably contributing to the incomplete assembly, as no long-range lattices can be formed.

To further understand the structural plasticity of the formed chimeric lattice, we performed a 3D variability analysis (3DVA)^29^ of the CH1 CLP SPA data. This showed that the lattice continuously samples the conformational space between curved and flat lattice, implying a tug-of-war between the two domains with both trying to satisfy their preferred geometry (Movies S1-3). To compare hexamer stability against overall lattice flexibility across the 3DVA space, RMSD values relative to wt were calculated for hexamer and 18-mer clusters (Table S3). Across all components, the hexamers remained highly stable for both HTLV-1 CA-NTD (ranging 2.0–2.3 Å) and HIV-1 CA-CTD-SP1 (1.1 Å). Notably, the CA-CTD-SP1 18-mer RMSD (1.7–2.1 Å) remained within the same range as the CA-NTD hexamer alone, indicating minimal additional flexibility contributed by the CA-CTD-SP1 dimeric inter-hexamer interface. In contrast, the CA-NTD 18-mer exhibited an increase in RMSD (3.0–3.5 Å), substantially higher than that of its own hexameric subunit and of the CA-CTD 18-mer. This implicates the CA-NTD inter-hexamer interfaces as the primary site of lattice flexibility. In contrast, both CA-NTD and CA-CTD-SP1 hexamers, along with the CA-CTD dimeric inter-hexamer interface, maintain a relatively rigid arrangement. To further pinpoint movements within the trimer and dimer interfaces, we measured the individual displacement of each helix 4 contributing to the HTLV-1 CA-NTD trimer and helix 9 forming the HIV-1 CA-CTD dimer for all the extremes of the 3DVA (Table S4, S5). Again, displacements were measured as the average Cα shift of all remaining helices 4 or helices 9 for trimer and dimer interfaces, respectively, after aligning one shared reference helix between wt and CH1 at each trimer or dimer interface. This showed that helix 9 (dimer interface) exhibits minimal displacement. The trimer interface instead displays a substantial displacement and a clear intra-interface asymmetry: within an 18-mer the six trimer interfaces can change irregularly, with helices 4 moving unevenly in vertical or horizontal directions (Fig. 1E) acting as a dynamic hinge that expands, retracts, and tilts to accommodate local lattice curvature.

### Linker length modulates assembly morphology and geometric tension of CH1

Despite the increased observed inter-domain distance and lack of interactions between CA-NTD and CA-CTD in CH1 (Fig. 1C), there appeared to be a communication between the domains in the form of a geometric structural tension leading to irregular CLP assemblies. Based on this observation we were wondering whether alleviating the tension between the two domains can result in more regular assemblies while satisfying geometric preferences of each domain.

To this end we designed a chimera with increased linker length between the CA domains. Specifically, we additionally inserted the HTLV-1 linker and 3_10_-helix segment (DPSWA) into CH1. This double linker (dl) CH1dl chimera also assembled into CLPs in HIV-1 immature assembly conditions (Fig. 2A). Interestingly, these CLPs appeared more complete than the CH1 CLPs, being oval in shape, with more curved lattice segments (55%) compared to flat/tubular lattice segments (20%). This suggests that extending the linker allows accommodating a stronger preference for uniform curvature which in turn alleviates the geometric strain that otherwise leads to broken assemblies.

**Figure 2:**
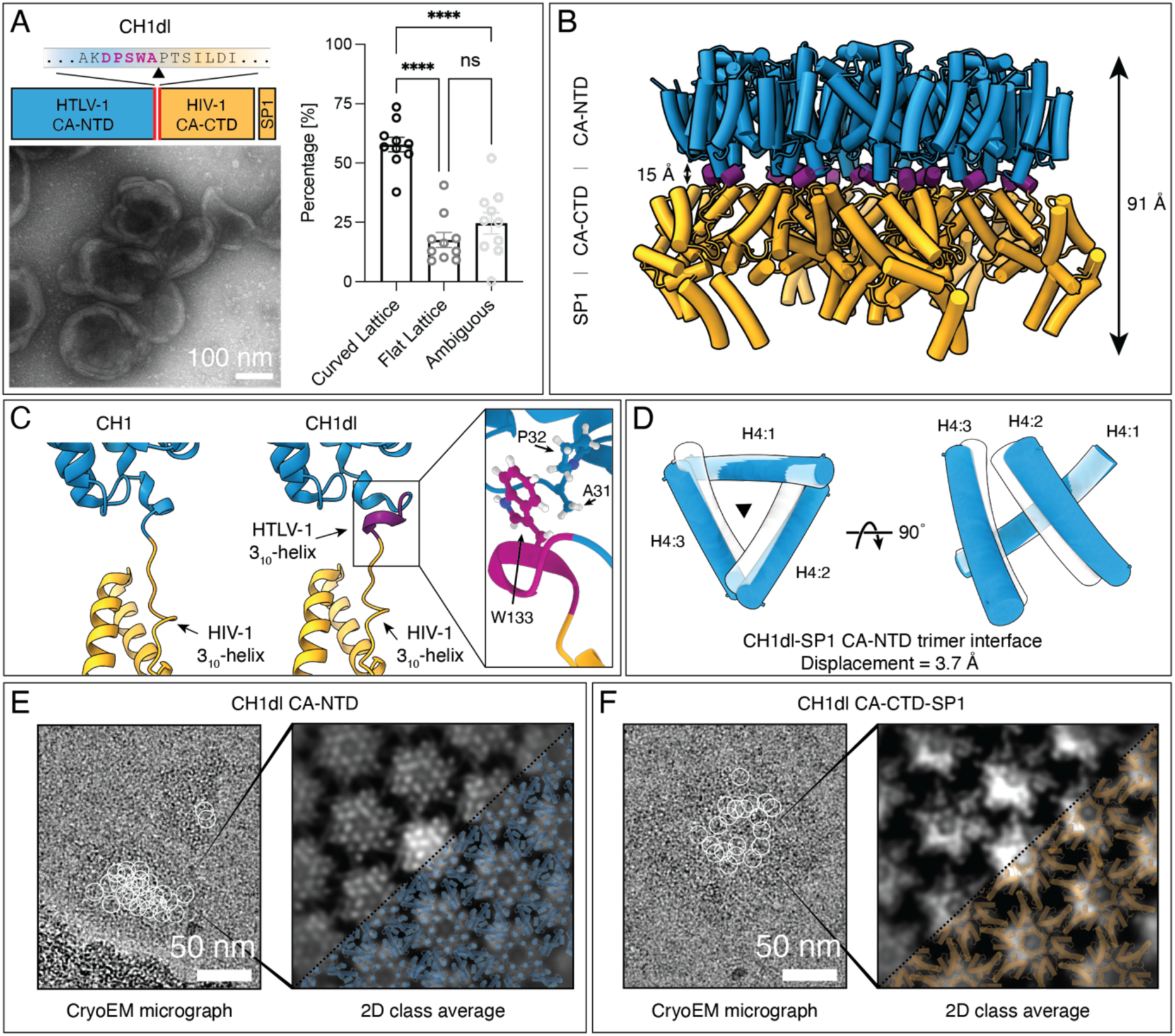
Structure of CH1dl. **A)** Top: Schematic of the CH1dl construct displaying the inserted HTLV-1 linker residues in purple. Bottom: Representative negative stain micrograph for CH1dl CLP assemblies. CLPs have a mean diameter of 195.16 nm (± 5.11 standard error of the mean (SEM)) and 143.64 nm (± 3.91) along the major and minor axes, and an aspect ratio of 1.39 ± 0.04 (mean ± SEM). Right: quantification of CLP morphology, **** p < 0.0001. **B)** Side view of the modeled CH1dl lattice, which displays similar height and inter-domain distances (91 Å and 15 Å, respectively) as CH1. The coloring is identical to Fig.1, with the HTLV-1 3_10_-helix colored in purple. **C)** Comparison of the linker of CH1 and CH1dl shows the location of the additional 3_10_-helix in CH1dl. The enlarged inset highlights the sequestering of the 3_10_-helix by the CA-NTD where W133 interacts with A31 and P32. The coloring is identical to B. **D)** The inter-hexamer CA-NTD trimer interface displacement in CH1dl compared to wt HTLV-1 CA-NTD (PDB 9N92). The coloring is identical to Fig. 1. **E-F)** Flat lattice sheets of CA-NTD (E) and CA-CTD-SP1 (F) as identified by particle picking (left) and after 2D-classification in cryoSPARC (right). White circles in the micrographs annotate particle picks that generated the respective 2D class averages. The models of either the HTLV-1 CA-NTD (blue) or the HIV-1 CA-CTD-SP1 (orange) are projected onto the 2D class averages, showing the good correspondence between class average and model and the lack of the density for the other domain. Scale bars for panels (A, E, F) are annotated in the figure.

We again performed SPA cryo-EM to determine the molecular interactions formed within CH1dl CLPs (Fig. S3, Table S2). The obtained 3.7 Å resolution structure revealed the same overall structural arrangement of both domains when compared to CH1 and an almost identical lattice height (91 Å), with a slightly increased CA-NTD/CA-CTD distance (15 Å) (Fig. 2B, Fig. S4A,B). CH1dl showed a weaker linker density than CH1, suggesting a higher degree of flexibility (Fig. 2C, Fig. S4C). However, modeling revealed the formation of two 3_10_-helices. While the positioning of the HIV-1 3_10_-helix was similar as in its wt context by binding the CA-CTD and contributing to the dimeric inter-hexameric interface, the HTLV-1 segment of the linker adopted a different positioning, due to the lack of its native binding site^10^. In CH1dl, the HTLV-1 linker segment is instead sequestered by the CA-NTD through a hydrophobic interaction between W133 and residues A31 and P32, at the base of helix 2 (Figure 2C). The CH1dl CA-NTD trimer interfaces dilated even more when compared to wt HTLV-1 (trimer interface displacement of 3.7 Å) (Fig. 2D). This finding implies that the non-native binding of the HTLV-1 3_10_-helix to the CA-NTD and the increased linker length promote both a stronger dilation/deformation of the trimeric interface and a more favorable setting for the CA-CTD-SP1 interactions to adopt their preferred curvature. As this can result in more long-range lattices, the CLPs appear more complete than the single-linker chimeras.

Interestingly, during 2D classification of the CH1dl cryo-EM data, we observed two distinct 2D classes, originating from flat lattice sheets not observed for CH1. No structure could be obtained because of the lack of side views, but by projecting the model of CH1dl onto these flat sheet 2D class averages, we observed that the classes correspond to either only HTLV-1 CA-NTD or only HIV-1 CA-CTD-SP1 (Fig. 2E,F). The flat lattice sheets may therefore represent assemblies where one subdomain fails to lock into its native arrangement.

### Lattice interface mutations and SP1 removal reveal independent and preferential assembly capabilities of CA-NTD and CA-CTD-SP1

After finding the CH1 subdomains can both form immature contacts outside of their native context, we aimed to further characterize their interdependency and their preferential geometries. We therefore created CH1 mutants that aimed at abolishing the assembly capabilities of either domain.

First, we mutated residues Y61 and H72 in the HTLV-1 CA-NTD which we recently described to be essential to stabilize intra-hexamer and inter-hexamer interfaces in the immature lattice^9^. The resulting mutant, termed CH1-NTDm failed to assemble in HTLV-1 assembly conditions, but intriguingly, assembled efficiently in HIV-1 assembly conditions where it adopted a regular icosahedral-like morphology (Fig. 3A), with particles having diameters of 58 ± 0.6 nm (mean ± SEM) and an increased layer thickness (∼20nm), as evidenced by nsTEM. To further characterize these particles and understand the dramatic shape change upon disrupting the CA-NTD interactions, we obtained a low-resolution (∼8 Å) SPA structure of this mutant (Figure S5). The structure revealed an unstructured HTLV-1 CA-NTD, while the HIV-1 CA-CTD-SP1 again adopted its native wt arrangement (Fig. S6). This shows that under permissive conditions (e.g., without a strong competing assembly domain) the CA-CTD can adopt a particle morphology with a high curvature. Interestingly, the size and morphology is very similar to particles formed by retrotransposons such as Ty1/Copia, Ty3/Gypsy or the capsid core of spumaviruses^30–32^.

**Figure 3:**
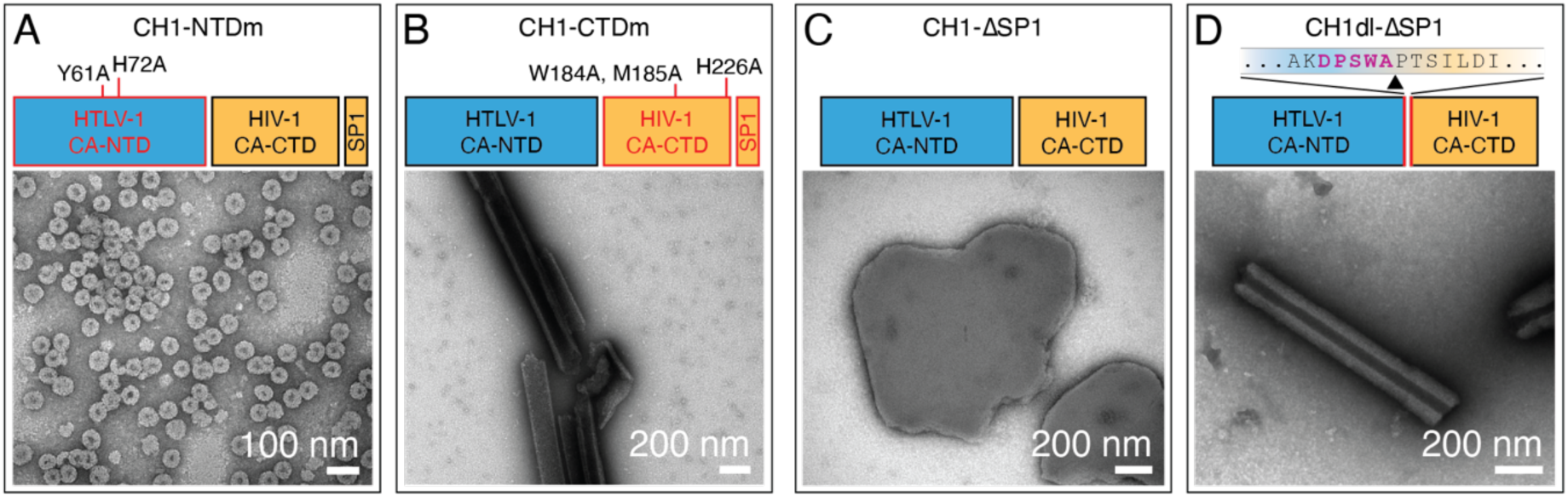
Chimera mutant assemblies. **A)** Schematic of the CH1-NTDm construct highlighting the mutated domain (CA-NTD, red) where the mutated residues are annotated. Below a representative negative stain TEM micrograph for CH1-NTDm CLP assemblies in HIV-1 immature assembly conditions is shown. **B)** Schematic of the CH1-CTDm construct highlighting the mutated domain (CA-CTD, red), where again the mutated residues are annotated. A representative negative stain TEM micrograph for CH1-CTDm CLP assemblies in HTLV-1 immature assembly conditions is shown below. **C)** Schematic of the CH1-ΔSP1construct with a representative negative stain TEM micrograph for CH1-ΔSP1 CLP assemblies in HIV-1 mature assembly conditions. **D)** Schematic of the CH1dl-ΔSP1construct, with a representative negative stain TEM micrograph for CH1dl-ΔSP1 CLP assemblies in HIV-1 mature assembly conditions. Adding the HTLV-1 linker segment to the ΔSP1 mutant shifts the morphology from flat lattice sheets to multi-layered tubular CLPs. Scale bars are annotated in all figure panels.

We next mutated residues in the C-terminal region of CH1, specifically W184, M185 (inter-hexamer), and H226 (intra-hexamer) in the HIV-1 CA-CTD to obtain a mutant termed CH1-CTDm. Again, as expected for mutating HIV-1 contacts, CH1-CTDm failed to assemble in HIV-1 conditions. Instead, under HTLV-1 assembly conditions, the CH1-CTDm formed tubular CLPs, albeit with very low efficiency and reproducibility (Fig. 3B). This shows that by providing the HTLV-1 CA-NTD with permissive conditions without a competitive assembly partner, it will revert back to an HTLV-1-like phenotype (see Fig. S1B). However, the low assembly efficiency also implies that a proper dimerizing CA-CTD domain is beneficial to potentially nucleate and stabilize the assembly. Results where the HTLV-1 CA-CTD-dimer interface has been mutated support this notion^10^.

Finally, we wanted to know if the assembly competence of chimeras is retained if HIV-1 SP1 is absent and if locking the CA-CTD in an immature state is essential for assembly driving capacity in the chimeric context. We designed two CH1 mutants (CH1-ΔSP1 and CH1dl-ΔSP1) and again subjected them to the same immature assembly conditions as for the previous constructs, where neither of them assembled. Since the ΔSP1 constructs did not assemble in immature assembly conditions, we decided to test both ΔSP1 mutants in conditions that have previously yielded mature capsid cores for HIV-1 CA^33^ (Table S1). In this setting, CH1-ΔSP1 formed predominantly flat lattice sheets (Fig. 3C), suggesting that the HTLV-1 CA-NTD drives an assembly similar to what we observed in 2D class averages of CH1dl (Fig. 2E). In the same conditions, CH1dl-ΔSP1 instead appeared as tubular CLP assemblies with thick multilayered walls (Fig. 3D). Hence, removal of SP1 still allows efficient but increasingly diverse and potentially non-immature chimeric assembly.

In summary, these mutants confirm that HTLV-1 CA-NTD and HIV-1 CA-CTD-SP1 are assembly drivers and nucleators able to kickstart their own assembly, albeit with vastly different efficiencies, and that their assembly preferences with respect to geometry and architecture are deeply encoded within the individual domains.

## Discussion

Here we have shown that CA subdomains from two evolutionary divergent retrovirus species, i.e., HTLV-1 CA-NTD and HIV-1 CA-CTD in conjunction with SP1, can assemble in vitro into their respective authentic immature assembly geometries even in a heterologous chimeric context. These data provide insight into the structural and architectural principles encoded into retroviral CA subdomains. Specifically, they illustrate the divergent evolutionary solutions of HIV-1 and HTLV-1 to accomplish the same architectural requirement of assembling metastable immature CA lattices, which remain predisposed to eventually converge onto identical mature CA lattices.

### Curvature and assembly morphology preferences are imprinted into retroviral CA subdomains

Our structural and mutagenesis data shows that both assembly competent CA subdomains of HTLV-1 and HIV-1 can drive assembly of an immature-like lattice independently of each other, but that they do prefer having a partner to initiate and stabilize CLP formation. However, our data also shows that this partner does not necessarily need to be properly structured, as evidenced from our NTDm and CTDm mutants and in line with our observations that the CA-CTD lattice in HTLV-1 VLPs and CLPs shows increased flexibility^9^. This is also reminiscent of early cryo-ET studies of the xenotropic murine leukemia virus-related virus (XMRV), where VLPs displayed an ordered CA-CTD lattice and a seemingly disordered CA-NTD^34^, suggesting that also in this retrovirus species the CA-CTD might be the assembly competent domain.

However, when heterologous assembly-competent subdomains are paired, a tug-of-war between them results in a strained lattice that gives rise to a mixed assembly phenotype. This reveals a distinct preference for adopting a flat lattice for HTLV-1 CA-NTD and curved lattice morphology for HIV-1 CA-SP1, respectively (Fig. 4). This is in line with previous observations that the HTLV-1 CA-NTD displays a strong preference to flat lattice areas within VLPs derived from cells expressing full-length Gag. It also agrees with earlier results describing the HIV-1 CA-CTD as minimal domain required for assembly and where the HIV-1 CA-NTD has been suggested to be more a modulator of shape, rather than assembly driver^35^. In the context of HTLV-1 the CA-CTD could take over a similar modulatory or at least lattice nucleation role, as it has been shown that the CA-CTD does not contribute strong interactions within the hexamer. In line with this, our data shows that CH2, where the assembly-passive CA-NTD of HIV-1 and CA-CTD of HTLV-1 are paired, is unable to generate an assembly-competent system under any of the tested conditions.

**Figure 4:**
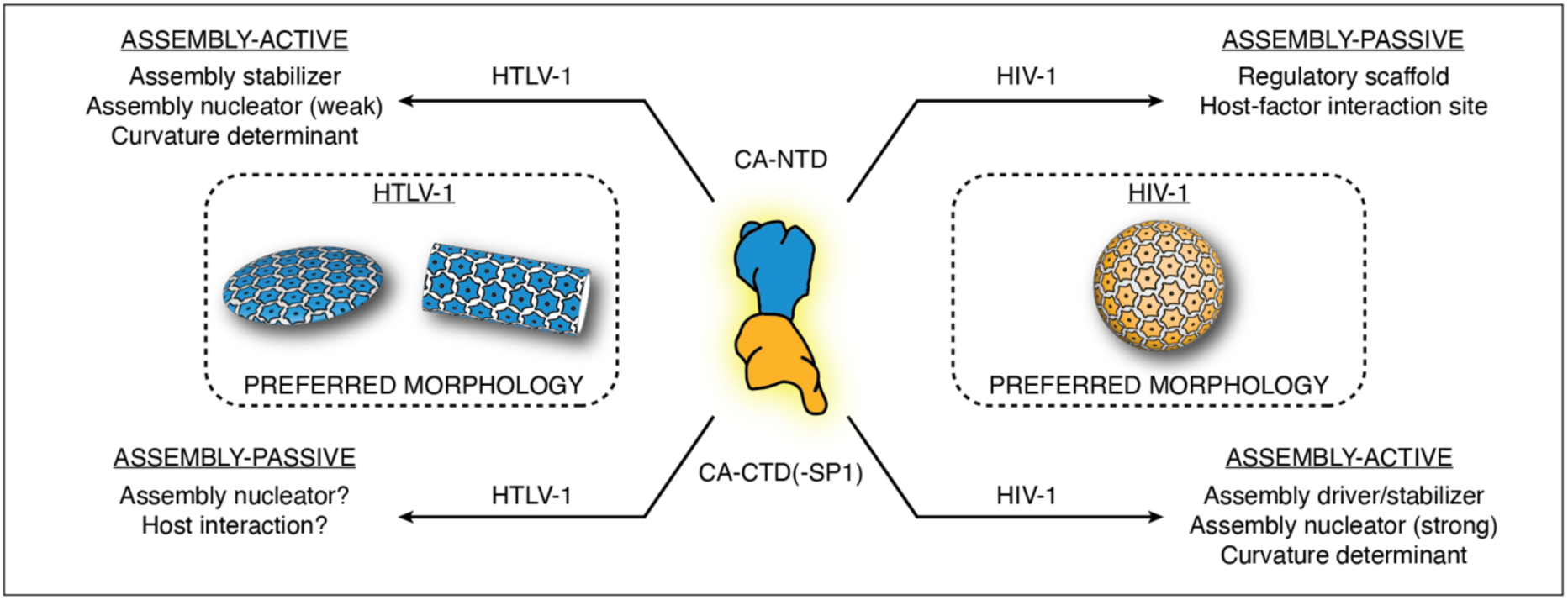
CA subdomain division of labor in immature assembly. Our chimeric and mutant CLP assemblies highlight that no strict co-evolution of CA subdomains is necessary for assembly, and that the two-domain capsid enables of a division of labor approach built on structural modularity. It appears that most of a retroviruses’ immature assembly properties can be encoded within a single CA-subdomain, resulting in immature assembly-passive and assembly active domains. The assembly-active HTLV-1 CA-NTD has a strong preference in forming flat lattice arrangements, while the assembly-active HIV-1 CA-CTD-SP1 is preferentially curved.

### Implications of geometrical CA subdomain preferences

Our data reveals further details about the extent of interface variation that makes the HTLV-1 CA-NTD still permissive to curvature, while trying to maintain a flat lattice. In HTLV-1, the CA-NTD hexamer represents a rigid building block, in line with what we already observed for in vitro assembled HTLV-1 MA_126_CANC tubes, where the hexamer barely deviated from 6-fold symmetry, despite assembling onto a two-fold symmetric tube^9^. Our 3DVA analysis clearly now shows the CA-NTD lattice to accommodate curvature variation through the dilation of its inter-hexamer trimer interface (Fig. 1E). Such plasticity at the trimeric CA-NTD interfaces and a strong rigidity of the hexamer within HTLV-1 is substantially different to observations of how the immature lattice in the lentivirus EIAV is stabilized, where the trimer represents the most stable contact point and variation and deformation occurs along dimeric inter-hexamer contacts and within the hexamer itself^8^.

The reasons for encoding the plasticity at the HTLV-1 NTD trimer interface may extend beyond curvature only. The HTLV-1 CA-NTD adopts a mature-like hexamer conformation even in the immature lattice^10,17^ and is thus hypothesized to undergo a less dramatic conformational change during maturation. Still, the immature CA-NTD contacts need to be broken to provide space for the mature CA-CTD rearrangement and an increased mature lattice spacing. Given the observed rigidity of the HTLV-1 CA-NTD hexamer, it is therefore plausible that the trimer interface acts as an intended lattice breaking point during maturation. Interestingly, studies have shown that a small binding pocket in the HTLV-1 CA-NTD can directly affect the trimer interface through conformational changes of residue H72^17^. This binding pocket is the equivalent site for binding CPSF6 and NUP153 in HIV-1 CA, and the potent HIV-1 capsid inhibitor Lenacapavir^36,37^. However, further experiments together with a structural determination of the mature lattice of HTLV-1 CA are needed to characterize the mechanisms releasing the trimer interactions.

In our chimera CH1, the CA-CTD-SP1 instead remains relatively unaltered across both the hexamer and inter-hexamer interfaces. By inserting additional residues in the linker of CH1 (CH1dl) the geometric tension between the two domains can be alleviated leading to more complete CLPs (Fig. 2A), within which the curvature dominates (HIV-1 CA-CTD-SP1 over HTLV-1 CA-NTD) and where the HTLV-1 CA-NTD inter-hexamer interface further dilates, demonstrating its curvature permissive nature.

### HTLV-1 displays an inverted distribution of assembly competence between the CA domains

The evolution of CA-NTD-driven assembly in HTLV-1 represents an alternative for achieving an immature assembly, in which the absence of strong CA-CTD-associated and downstream contacts permits the CA-NTD to assume the primary role in lattice stabilization. This potentially simpler assembly pathway is also reflected in full-length HTLV-1 Gag, which consists solely of the canonical MA, CA and NC domains and lacks a spacer peptide or other known assembly-stabilizing elements downstream of CA. Such an arrangement may result in a less pronounced structural distinction between the interaction networks stabilizing the immature and mature assemblies. This is consistent with the above-explained observation that the immature HTLV-1 CA-NTD lattice closely resembles the mature CA hexameric organization observed in other structurally characterized retroviruses.

It has been proposed that HIV-1 maturation can involve a displacive transition through an intermediate in which the HIV-1 CA-NTD adopts a mature hexameric lattice configuration with immature lattice spacing, while the CA-CTD–SP1 region retains its immature-like form^38^. Since the immature HTLV-1 CA-NTD hexamer in CH1 is similar to a mature HIV-1 CA-NTD hexamer, while being combined with an immature HIV-1 CA-CTD-SP1, our CH1 construct replicates the geometry of that hybrid maturation intermediate. The natural occurrence of a comparable hybrid organization during HIV-1 maturation provides a structural precedent for our chimeric assembly and may help explain how the HIV-1 CA-CTD–SP1 hexamer can accommodate a substantially altered CA-NTD lattice architecture. Our findings further suggest that native, species-matched interactions between the CA-NTD and CA-CTD are not strictly required to establish such a hybrid lattice.

The adaptation of HTLV-1 to rely predominantly on the CA-NTD for immature assembly is particularly striking given the stronger evolutionary conservation of the CA-CTD, both in sequence and protein fold. In most structurally characterized retroviruses, the CA-CTD provides conserved interactions important for immature lattice formation and hexamer stabilization, often together with regions immediately downstream of CA. Interestingly, our CH1-NTDm construct yields small, regular particles that are morphologically reminiscent of capsid-like assemblies formed by retrotransposons such as Ty1 and Ty3/Gypsy^30,32^ (Fig. 3A). The two-domain CA architecture itself is thought to have arisen through an ancient duplication of a CA-like fold, after which the CA-NTD diverged more extensively than the CA-CTD. This raises the possibility that, under permissive conditions, the HIV-1 CA-CTD–SP1 region retains the capacity to drive a simpler mode of assembly when paired with an essentially non-functional HTLV-1 CA-NTD.

More generally, the stronger conservation of CA-CTD-mediated assembly may have allowed the CA-NTD of many retroviral lineages to diversify without necessarily abolishing particle formation. Such freedom could have facilitated the acquisition of different CA-NTD-mediated interactions and assembly geometries, as observed in structural studies of immature CA lattices in different retroviral species^5–9^. The retroviral CA-NTD contributes stabilizing contacts in both immature and mature lattices, but also occupies a more exposed position and participates in interactions with cellular factors, as demonstrated for HIV-1^36,39–41^. This broader functional and structural exposure may have contributed to its greater evolutionary divergence.

From this perspective, the switch of HTLV-1 towards CA-NTD-driven immature assembly is particularly interesting. For the CA-NTD to become the dominant assembly determinant, assembly-promoting CA-NTD contacts must have become sufficiently strong while the relative contribution of the CA-CTD to lattice organization was reduced. These changes need not have occurred simultaneously; intermediate states in which the CA-NTD and CA-CTD shared assembly competence are also conceivable. Whether such a balance exists in other retroviruses with divergent immature CA quaternary arrangements represents an interesting target for future investigation.

An important feature in our CH1-NTDm construct is the presence of HIV-1 SP1. Because the contribution of SP1 to lattice organization and curvature in this altered assembly context is unclear, it would be informative to characterize CH1-NTDm variants lacking SP1. Similarly, chimeras combining CA domains from retrotransposons with those of HIV-1 or HTLV-1 could be used to test assembly compatibility across a substantially greater evolutionary distance. Finally, determining whether the HTLV-1 CA-NTD has similarly extensive interactions with host factors could help establish whether its unusual role in immature assembly is accompanied by broader functional constraints on CA-NTD evolution.

In summary, the chimera system presented here provides a unique experimental framework to comparatively study assembly properties of CA subdomains. We have shown that retroviral CA subdomains encode autonomous and competing geometric preferences that have evolved independently, which has broad implications for understanding how assembly diversity has been generated across retroviral species.

## Material and Methods

### Cloning, expression, and purification of HTLV-1 CA, HIV-1 CA-SP1, and chimeras

Codon-optimized sequences of HTLV-1 CA (Gag residues 131-344, UniProt code P03345), HIV-1 CA-SP1 (133-377, UniProt code P04585), CH1 (131-259 of HTLV-1 Gag and 279-377 of HIV-1 Gag), and CH2 (131-278 of HIV-1 Gag and 260-344 of HTLV-1 Gag) with a C-terminally placed 6xHis-tag were separately cloned into pET28 expression vectors encoding kanamycin resistance, downstream of a 6xHis-SUMO tag, using standard molecular cloning methods. All mutants were created by around-the-horn PCR to edit required nucleotides. The vectors were transformed into competent Escherichia coli BL21 cells and grown overnight in lysogeny broth (LB) medium supplemented with kanamycin at 37 °C and 220 rpm. For all constructs except CH1-ΔSP1, the overnight culture was used to inoculate 1-2 L of LB medium at a 1:20 ratio, and the cells were allowed to grow at 37 °C and 220 rpm until an optical density at 600 nm (OD600) of 0.5-0.7 was reached. Protein expression was then induced by adding IPTG at a final concentration of 1 mM. The cells were harvested after 5-6 h induction at 37 °C and 200 rpm, by centrifugation at 6,000g for 15 min. For CH1-ΔSP1, the cells were instead harvested directly after overnight incubation. The resulting pellets were resuspended in 20 mM Tris, 500 mM NaCl, 2 µM ZnCl_2_, 5% glycerol, 1 mM PMSF and 1 mM TCEP, pH 8 (resuspension buffer). To extract the proteins, the cells were lysed via 3-5 freeze-thaw cycles at -80 °C and 42 °C respectively. Cell debris was pelleted by centrifugation for 1 h at 21,000g and 4 °C. The resulting supernatant containing the protein was subjected to nucleic acid precipitation using polyethylenimine (PEI) at a final concentration of 0.3 % under stirring at 4 °C for 15 min. Nucleic acid was disposed of by centrifugation at 10,000g, and the supernatant containing the protein was subjected to ammonium sulfate protein precipitation at a 40 % saturation under stirring at 4 °C overnight. The protein was collected by centrifugation at 10,000g at 4 °C and resuspended in 20 mM Tris, 20 mM imidazole, 500 mM NaCl, and 0.5 mM TCEP, pH 8 (wash buffer). The proteins were then purified using affinity chromatography with 1 mL HisTrap FF columns (Cytiva, 17531901) on a Cytiva AEKTA pure according to the manufacturers’ instructions and eluted using 20 mM TRIS, 500 mM imidazole, 500 mM NaCl, and 0.5 mM TCEP, pH 8 (elution buffer). After affinity chromatography, the 6xHis-SUMO tag was cleaved by addition of Ulp1 during dialysis into 20 mM TRIS, 500 mM NaCl, and 0.5 mM TCEP, pH 8 (storage buffer) overnight at 4 °C. Because of the presence of a BamHI cloning site between the 6xHis-SUMO and the target construct, all cleaved target proteins have an ectopic serine and the very N-terminus. The proteins were then separated from the cleaved tag by size exclusion chromatography using a HiLoad 26/600 Superdex 75 pg column (Cytiva, 28989334) and storage buffer on a Cytiva AEKTA pure according to the manufacturers’ instructions. The purified proteins were flash-frozen in liquid nitrogen at 5 mg/ml in 400 µL aliquots and stored at -80 °C.

### In vitro assembly and morphology quantification

In vitro assembly of all constructs was done in two different immature assembly conditions established for either HTLV-1 or HIV-1^9,25^ (see Table S1). For HTLV-1 conditions, the proteins were concentrated to ∼680 µM and incubated in 50 mM MES and 200 mM NaCl, pH 6 at 4 °C for 4 h and then at 26 °C overnight. For HIV-1 immature conditions, proteins were concentrated to 500 µM and dialyzed into 50 mM MES, pH 6 with 500 µM IP6 and incubated for 2h. For mature HIV-1 conditions, proteins were concentrated to 100 µM and buffer-exchanged into 50 mM MES, 40 mM NaCl and 1 mM TCEP, pH 6, followed by addition of 100 µM IP6 an incubation at 37 °C for 1h. Assemblies were visualized on a Tecnai T10 transmission EM instrument using negative staining with 2% uranyl acetate (UA).

To quantify CLP lattice morphologies, nsTEM micrographs were analyzed by a grid-based sampling in ImageJ. Grid intersections falling on CLPs were categorized as flat, curved, or ambiguous/obscured. For each construct, 10 micrographs across two independent replicates were evaluated for a total of 6,900 intersections per condition/construct. Analyses were done in GraphPad Prism (v11.1.0). Comparisons across groups used one-way ANOVA followed by Tukey’s post-hoc test. Error bars depict mean ± SEM.

### Vitrification of CH1, CH1dl, and CH1-NTDm CLPs

CH1, CH1dl, and CH1-NTDm CLP assemblies were vitrified using a Leica EM GP2 plunge-freezer. A 3.5 µL volume of assembly solution was deposited onto glow discharged (Quorum GloQube Plus, 1 min, 25 mA) 200-mesh holey carbon grids (Quantifoil R 2/2 X-103-Cu200) with the blotting chamber conditions set to 10 °C and 90 % humidity. Grids were blotted from the back for 5-6 s, then subsequently vitrified in liquid ethane (-185 °C) and stored under liquid nitrogen conditions until imaging.

### CH1 and CH1-NTDm cryo-electron microscopy data acquisition

Micrographs were acquired under cryogenic conditions on a Glacios cryo-TEM (Thermo Fisher Scientific) operating at 200 kV, equipped with an X-FEG electron source and a Falcon 4i direct electron detector. Data were recorded in EER format using EPU v3.9.1.8206 (Thermo Fisher Scientific) at a nominal magnification of 92,000x and 120,000x, corresponding to a pixel size of 1.511 Å and 1.182 Å for CH1 and CH1-NTDm, respectively. The total exposure dose was 50 e/Å² over 14.77 s at a dose rate of 3.38 e/Å²/s for CH1 and over 9.51 s at a dose rate of 5.25 e/Å²/s for CH1-NTDm. The nominal defocus range was set to −1.2 to −1.8 μm in 0.2 µm steps. A C2 aperture of 50 μm was used in nanoprobe mode with spot size 5 for CH1 and spot size 4 for CH1-NTDm.

### CH1dl-cryo-electron microscopy data acquisition

Micrographs were acquired under cryogenic conditions on a Titan Krios G3i cryo-TEM (Thermo Fisher Scientific) operating at 300 kV, equipped with an X-FEG electron source, a Gatan K3 direct electron detector and a BioQuantum post-column energy filter operated with a slit width of 20 eV. Automated data collection was performed using EPU 3.4.0 (Thermo Fisher Scientific) at a nominal magnification of 105,000x, corresponding to a pixel size of 1.089 Å. Movies were recorded in counting mode and fractionated into 48 frames over a 2.95 s exposure, with a dose rate of 32.86 e/pixel/s and a total accumulated dose of 80 e/Å². The nominal defocus was varied between −1.2 and −1.8 μm in 0.2 μm steps. A C2 aperture of 50 μm and an objective aperture of 100 μm were used at spot size 4.

### Cryo-EM data processing

Cryo-EM data processing was performed in cryoSPARC v4.6.2^42^ following a standardized workflow across constructs (Figs. S2, S3, S5). Raw movies were motion-corrected and dose-weighted using Patch Motion Correction, followed by contrast transfer function (CTF) estimation via Patch CTF Estimation. Automated particle picking was then performed using the Blob Picker with a search diameter set between 20 Å and 80 Å, and particles were initially extracted for downstream classification.

Multiple rounds of 2D classification were conducted to clear non-particle features and select clean CLP assemblies. For high-resolution datasets, selected 2D class averages were used to generate C1 ab initio reconstructions, followed by sequential rounds of C1 and C6 homogeneous refinement. Particle stacks were further pruned via C6 3D classification without alignment to remove bad particles. Optimal classes were subjected to iterative rounds of C6 non-uniform refinement, local refinement, local CTF estimation/refinement, and reference-based motion correction^43,44^. For the lower-resolution CH1-NTDm, clean 2D class averages were directly used to generate C6 ab initio reconstructions followed by C6 homogeneous refinement.

All final maps were locally filtered and adjusted for handedness. Specific dataset sizes, particle extraction parameters, box sizes, sub-classification breakdowns, and nominal resolutions for each construct are detailed in the respective processing schemes (Fig. S2, S3, S5) and summarized in Table S2.

### Modeling and RMSD measurements of CH1 and CH1dl

To model CH1 and CH1dl, existing structures of HTLV-1 CA-NTD (PDB 9N92) and HIV-1 CA-CTD-SP1 (PDB 5L93) were used as initial templates. The residues were manually fitted and adjusted in ISOLDE v1.11.1^45^ (integrated within UCSF ChimeraX v1.11.1^46^) to refine local stereochemistry. The resulting models were used to build the subunits at the inter-hexameric interface by rigid body fitting, and then C6 symmetry was applied to propagate the full 18-mer lattices. The 18-mers were then subjected to real-space refinements in Phenix v1.21.2^47^ using secondary structure and NCS restraints. All RMSD and distance measurements were calculated in ChimeraX v1.11.1 for backbone C*a* using the align command. Displacements of the trimer and dimer interfaces was assessed in ChimeraX by aligning one helix in the interface, then measuring the resulting average Cα displacement of all residues in the remaining helix/helices. All residue numbering in the chimeras is using the respective HTLV-1 CA-NTD and HIV-1 CA-CTD-SP1 numbers as shown in Figure S1A and S1B, respectively. 3DVA 18-mers were built by rigid-body fitting individual CA-NTD and CA-CTD domains across the full 18-mer lattices of each 3DVA components.

## Acknowledgements

We thank B. Arragain and other members of the Schur group for helpful discussion and support. FKMS acknowledges support from the Institute of Science and Technology Austria (ISTA). This research was supported by the Scientific Service Units (SSUs) of ISTA through resources provided by Scientific Computing (SciComp), the Life Science Facility (LSF), and the Electron Microscopy Facility (EMF).

## Contributions

Project administration, supervision, and funding acquisition: F.K.M.S.; conceptualization: M.P., R.A.D and F.K.M.S.; methodology: M.P. and F.K.M.S; investigation: M.P., A.T. and V.-V.H.; validation, formal analysis, and visualization: M.P. and F.K.M.S.; data curation: M.P.; writing – original draft: M.P. and F.K.M.S.; writing – review and editing: M.P., A.T., V.-V.H., R.A.D. and F.K.M.S.

## Competing Interests

The authors declare no competing interests.

## Data availability

The coordinates and EM maps generated in this study have been deposited in the Protein Data Bank and the Electron Microscopy Data Bank (see Table S2) under accession codes:

CH1: PDB 34IT EMD-59880

CH1dl: PDB 34IU EMD-59881

CH1-NTDm: EMD-59882

## Supplementary Figures

**Figure S1:**
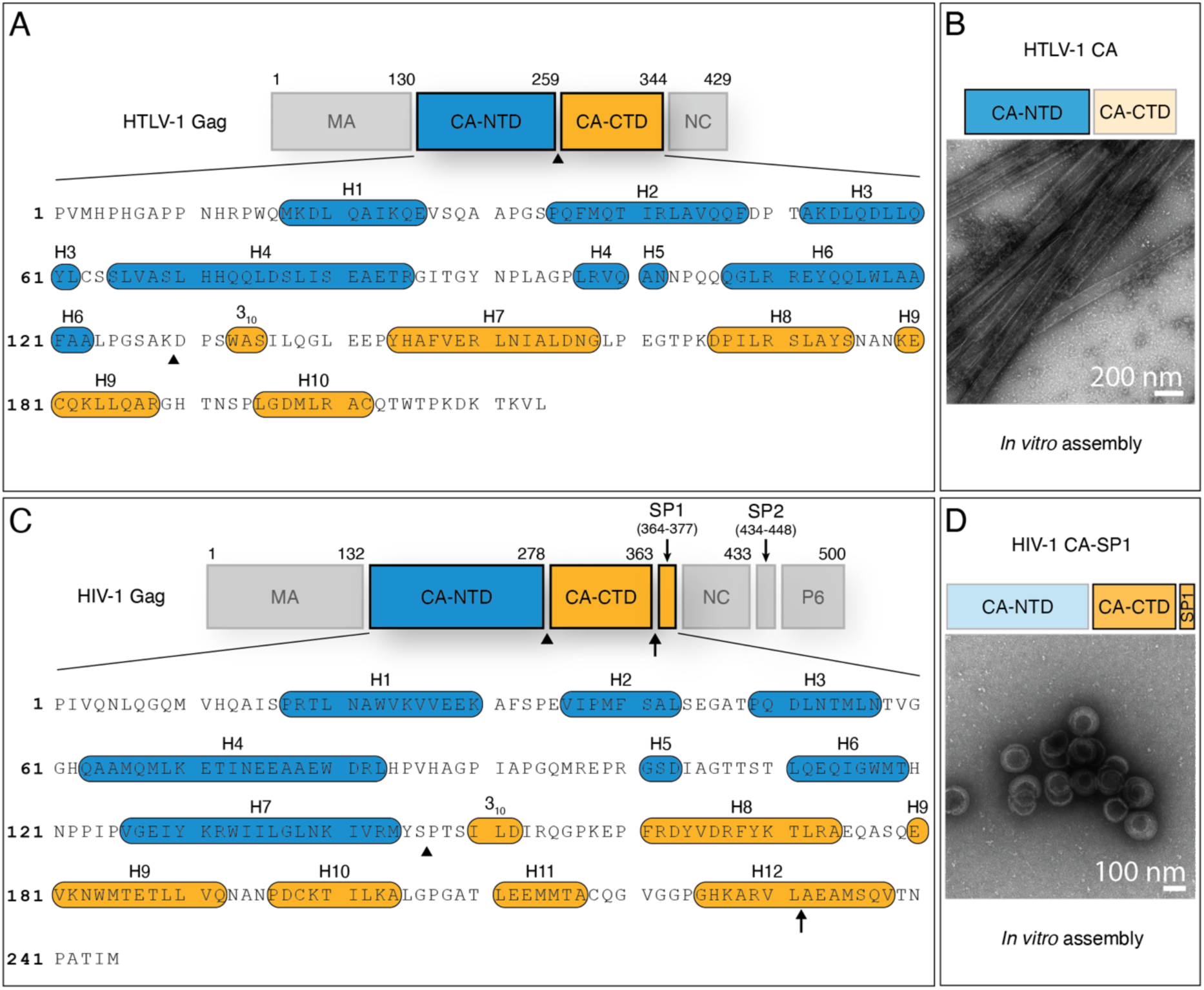
HTLV-1 and HIV-1 Gag, and *in vitro* CA assembly. **A.** HTLV-1 Gag schematic and CA sequence (UniProt code P03345). The CA-NTD (blue) and CA-CTD (orange) are highlighted. CA-sequence numbering starts from 1. Alpha-helical secondary structures are highlighted. The boundary between the CA subdomains (residues K129 and D130) is annotated with a black arrowhead. **B.** Representative negative stain TEM micrograph for HTLV-1 CA CLP assemblies. **C.** HIV-1 Gag schematic and CA sequence (UniProt code P04585). Coloring and annotations are identical to A. The boundary between CA subdomains (residues S147 and P148) is annotated with a black arrowhead. The cleavage site between HIV-1 CA-CTD and SP1 (L231 and A232) is annotated with a black arrow. **D.** Representative negative stain micrograph for HIV-1 CA-SP1 CLP assemblies. Scale bars for panels B and D are annotated within the figure. Assembly conditions for HIV-1 CA-SP1 and HTLV-1 CA are given in Table S1.

**Figure S2:**
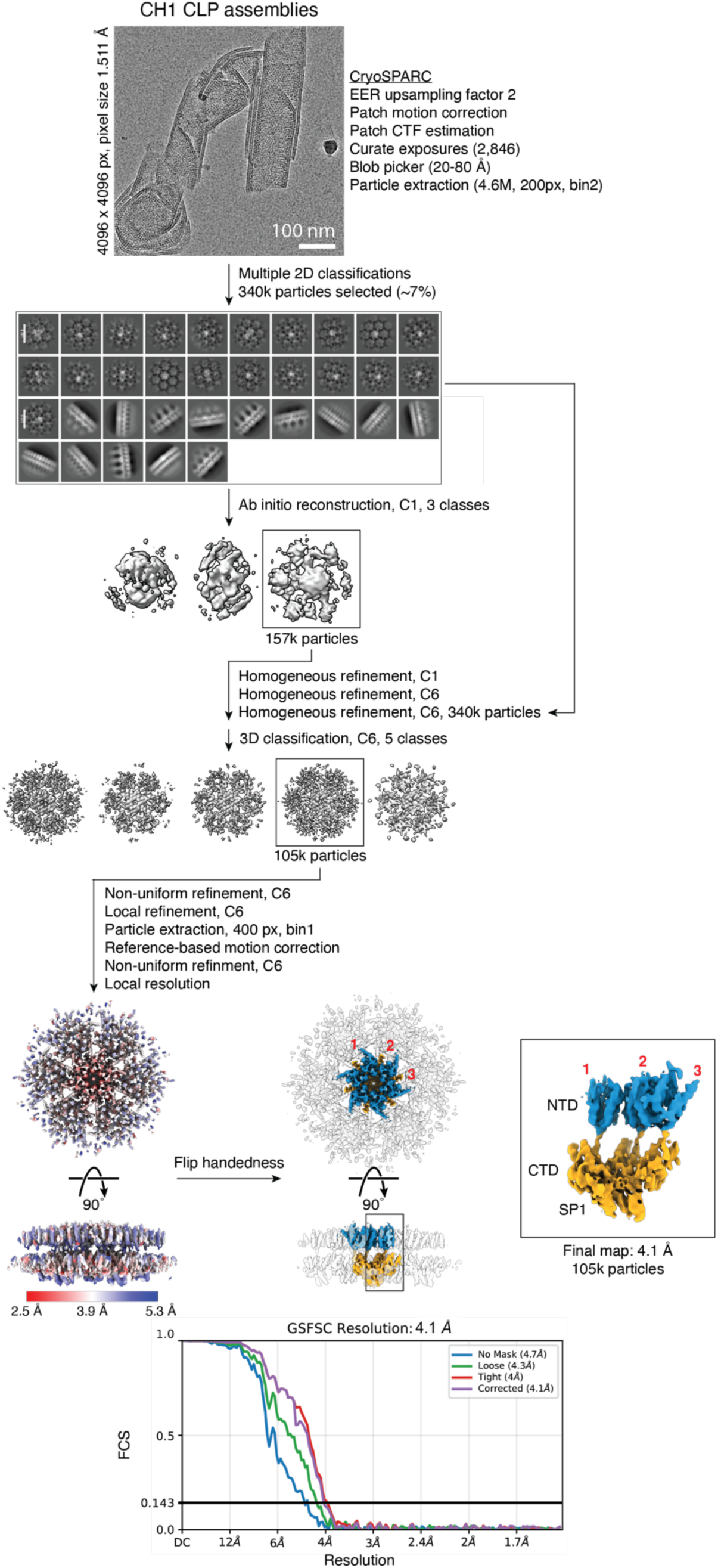
Cryo-EM data processing workflow of CH1. The entire data processing pipeline was performed in cryoSPARC v4.6.2. High-quality particles were selected through multiple 2D classification rounds and used for ab-initio reconstruction with no symmetry imposed. The resulting best structure was subjected to multiple rounds of homogeneous refinement without and with six-fold symmetry, followed by 3D classification to select for the best particle class, before being further refined. The final reconstruction shows the reconstructed lattice with a hexamer colored in blue (CA-NTD) and orange (CA-CTD-SP1).

**Figure S3:**
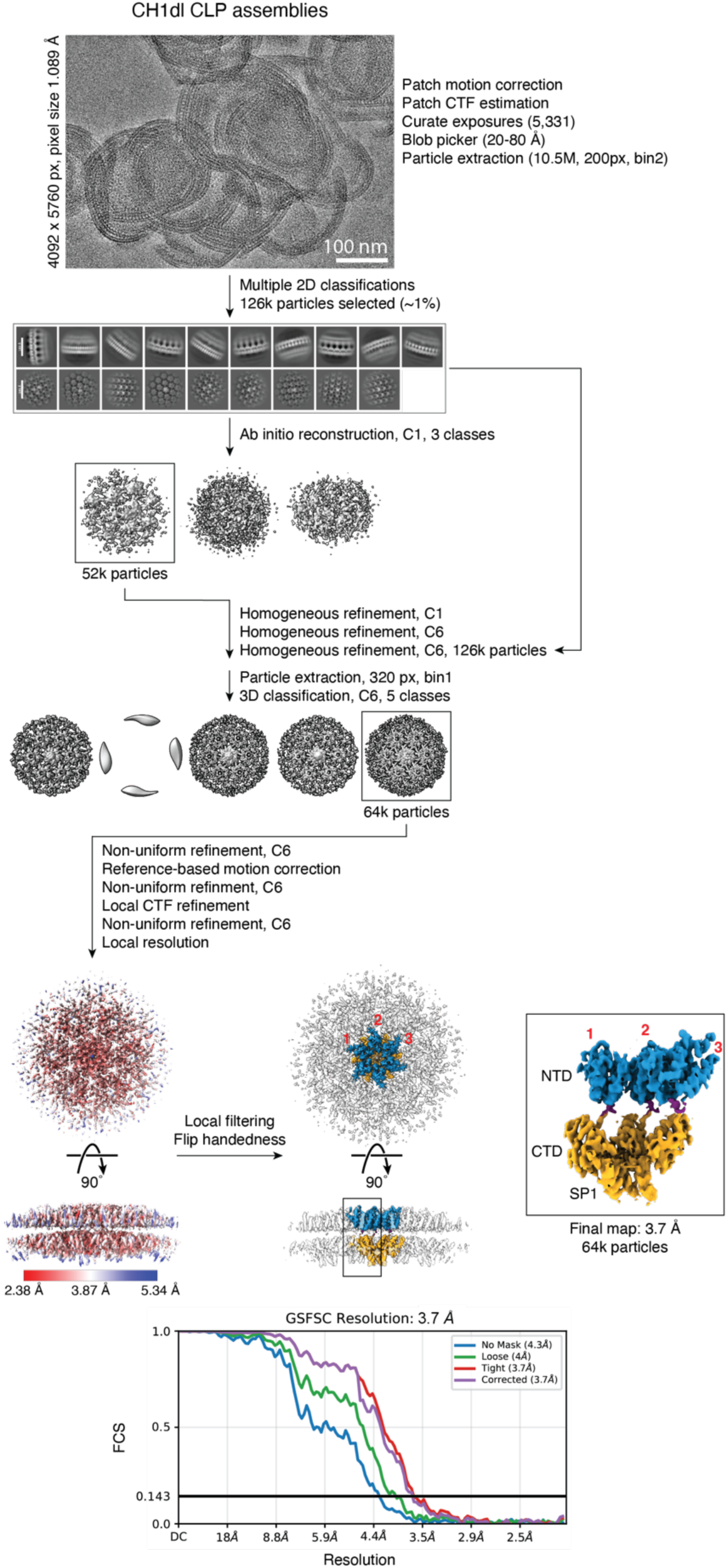
Cryo-EM data processing workflow of CH1dl. The entire data processing pipeline was performed in cryoSPARC v4.6.2. High-quality particles were selected through multiple 2D classification rounds and used for ab-initio reconstruction with no symmetry imposed. The resulting best structure was subjected to multiple rounds of homogeneous refinement without and with six-fold symmetry, followed by 3D classification to select for the best particle class, before being further refined. The final reconstruction is colored identically to Fig. S1.

**Figure S4:**
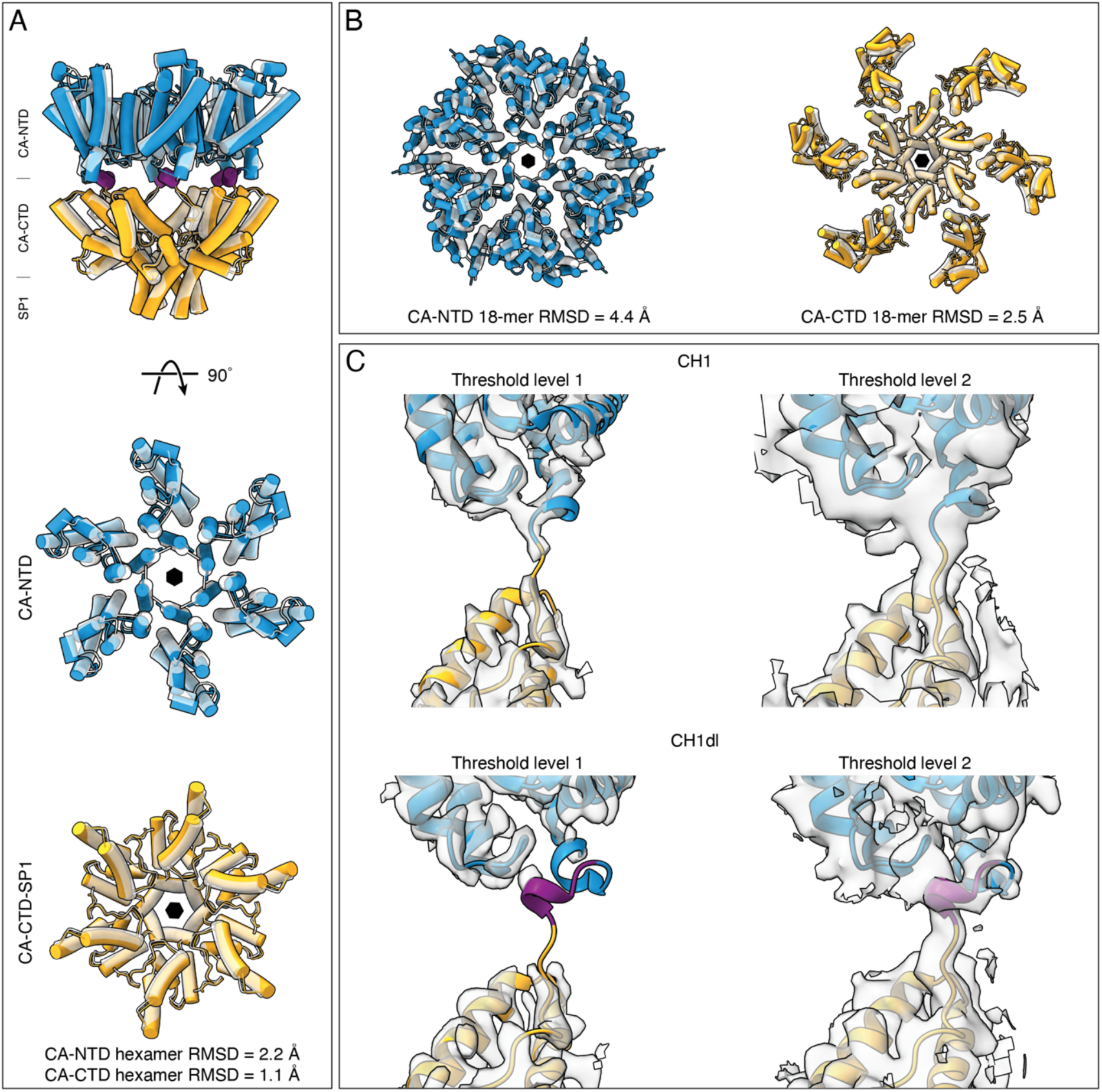
Comparative analysis of CH1dl. **A)** Alignment of a hexamer of CH1dl with hexamers of HTLV-1 CA-NTD (PDB 9N92, RMSD 2.2 Å) and HIV-1 CA-CTD SP1 (PDB 5L93, RMSD 1.1 Å). Coloring is identical to Figure 2. **B)** Alignment of an 18-mer of CH1dl with 18-mers of HTLV-1 CA-NTD (RMSD 4.4 Å) and HIV-1 CA-CTD-SP1 (RMSD 2.5 Å). **C)** Comparison of linker densities from the SPA maps of CH1 and CH1dl at two different threshold levels, demonstrating an overall weaker linker density in CH1dl, but a clear density for the HTLV-1 3_10_-helix connecting to the base of the HTLV-1 CA-NTD.

**Figure S5:**
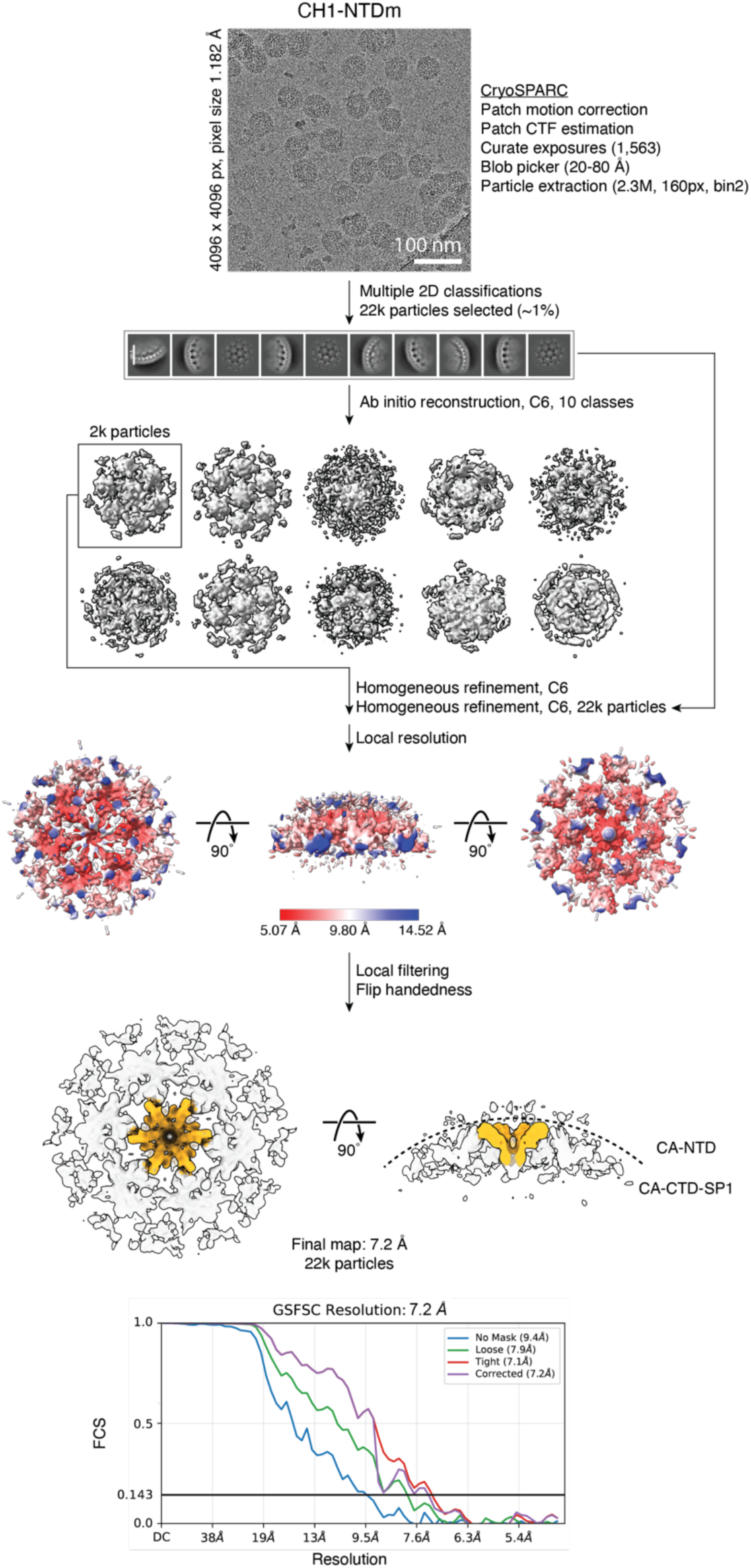
Cryo-EM data processing workflow of CH1-NTDm. The entire data processing pipeline was performed in cryoSPARC v4.6.2. High-quality particles were selected through multiple 2D classification rounds and used for ab-initio reconstruction with C6 symmetry imposed. The resulting best structure was subjected to multiple rounds of homogeneous refinement. The final reconstruction is colored identically to Fig. S1.

**Figure S6:**
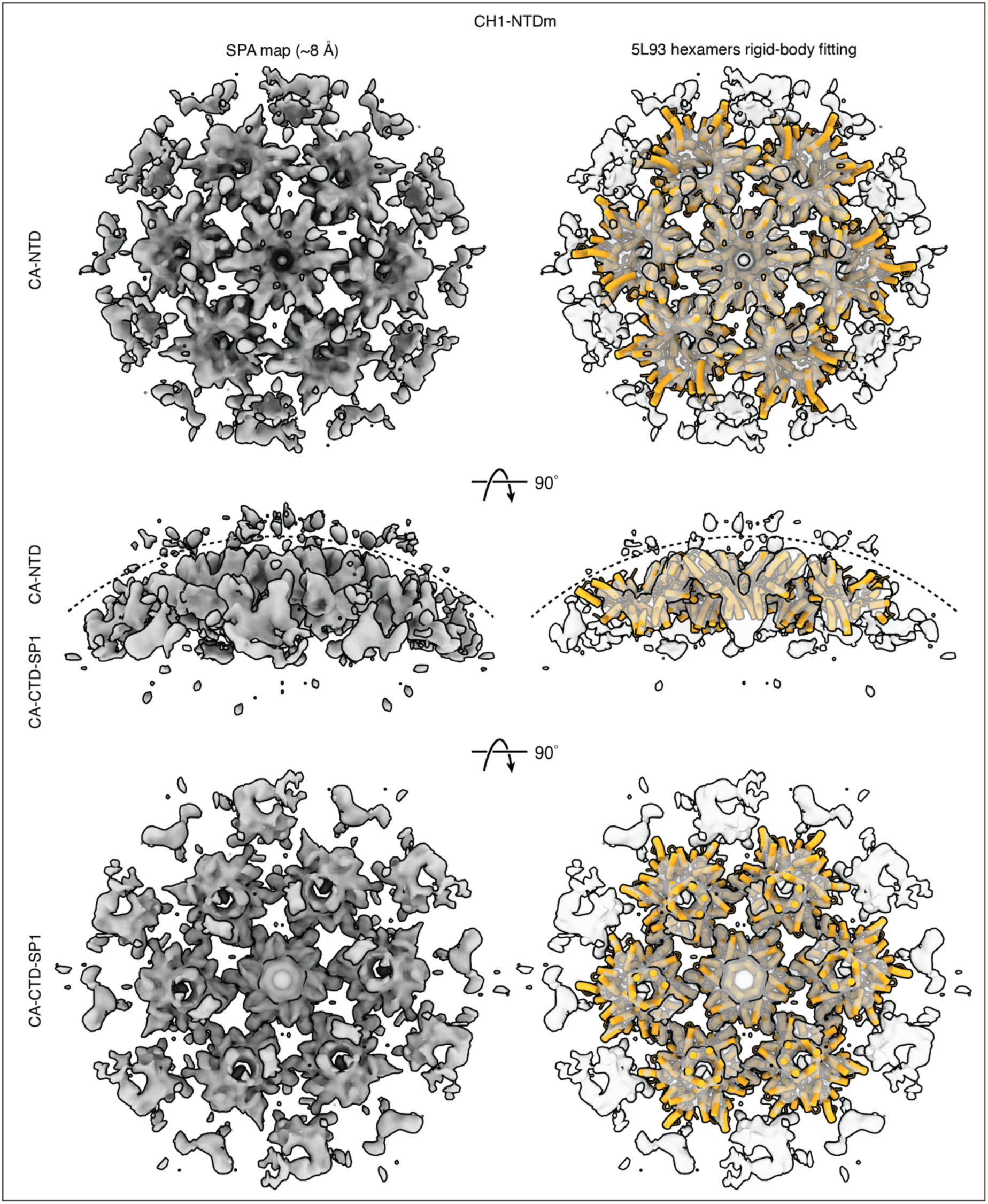
Immature CA-CTD lattice and unresolved CA-NTD in CH1-NTDm CLPs. Low-resolution (∼8Å) map views (top, side, and bottom) of CH1-NTDm without and with rigid-body fitted HIV-1 CA-CTD-SP1 (PDB 5L93) hexamers. The mutated HTLV-1 CA-NTD is not resolved. The fitted HIV-1 CA-CTD-SP1 is colored in orange.

**Movies S1–S3. 3DVA of lattice CH1 curvature**

Movies S1, S2, and S3 show 3DVA components 1, 2, and 3, respectively, each oscillating between its two extreme conformational states, illustrating the continuous sampling of curvature space by the chimeric CH1 CLP lattice. The CH1 lattice is shown from the top and two orthogonal views.

## Notes

### Competing Interest Statement

The authors have declared no competing interest.

## References

1. Accola, M. A., Höglund, S. & Göttlinger, H. G. A putative alpha-helical structure which overlaps the capsid-p2 boundary in the human immunodeficiency virus type 1 Gag precursor is crucial for viral particle assembly. J. Virol. 72, 2072–2078 (1998).

2. Sundquist, W. I. & Krausslich, H.-G. HIV-1 Assembly, Budding, and Maturation. Cold Spring Harb. Perspect. Med. 2, a006924–a006924 (2012).

3. Mattei, S., Schur, F. K. & Briggs, J. A. Retrovirus maturation-an extraordinary structural transformation. Curr. Opin. Virol. 18, 27–35 (2016).

4. Pornillos, O. & Ganser-Pornillos, B. K. Maturation of retroviruses. Curr. Opin. Virol. 36, 47–55 (2019).

5. Schur, F. K. M. et al. Structure of the immature HIV-1 capsid in intact virus particles at 8.8 Å resolution. Nature 517, 505–508 (2015).

6. Schur, F. K. M., Dick, R. A., Hagen, W. J. H., Vogt, V. M. & Briggs, J. A. G. The Structure of Immature Virus-Like Rous Sarcoma Virus Gag Particles Reveals a Structural Role for the p10 Domain in Assembly. J. Virol. 89, 10294–10302 (2015).

7. Qu, K. et al. Structure and architecture of immature and mature murine leukemia virus capsids. Proc. Natl. Acad. Sci. 115, (2018).

8. Dick, R. A. et al. Structures of immature EIAV Gag lattices reveal a conserved role for IP6 in lentivirus assembly. PLOS Pathog. 16, e1008277 (2020).

9. Obr, M. et al. Distinct stabilization of the human T cell leukemia virus type 1 immature Gag lattice. Nat. Struct. Mol. Biol. 32, 268–276 (2025).

10. Arndt, W. G. et al. High-resolution analysis of the human T-cell leukemia virus capsid protein reveals insights into immature particle morphology. Nat. Commun. 17, 444 (2025).

11. Bush, D. L. et al. Higher-order structure of the Rous sarcoma virus SP assembly domain. J. Virol. 88, 5617–5629 (2014).

12. Bharat, T. A. M. et al. Structure of the immature retroviral capsid at 8 Å resolution by cryo-electron microscopy. Nature 487, 385–389 (2012).

13. Cheslock, S. R. et al. Charged Assembly Helix Motif in Murine Leukemia Virus Capsid: an Important Region for Virus Assembly and Particle Size Determination. J. Virol. 77, 7058–7066 (2003).

14. Doležal, M. et al. Functional and Structural Characterization of Novel Type of Linker Connecting Capsid and Nucleocapsid Protein Domains in Murine Leukemia Virus. J. Biol. Chem. 291, 20630–20642 (2016).

15. Oshima, M. et al. Effects of blocking individual maturation cleavages in murine leukemia virus gag. J. Virol. 78, 1411–1420 (2004).

16. Maldonado, J., Cao, S., Zhang, W. & Mansky, L. Distinct Morphology of Human T-Cell Leukemia Virus Type 1-Like Particles. Viruses 8, 132 (2016).

17. Yu, R., Phalora, P., Li, N., Böcking, T. & Jacques, D. A. The Human T-cell Leukemia Virus capsid protein is a potential drug target. Nat. Commun. 16, 10892 (2025).

18. Taylor, W. R., Stoye, J. P. & Taylor, I. A. A comparative analysis of the foamy and ortho virus capsid structures reveals an ancient domain duplication. BMC Struct. Biol. 17, 3 (2017).

19. Martin, J. L. et al. Disparate Contributions of Human Retrovirus Capsid Subdomains to Gag-Gag Oligomerization, Virus Morphology, and Particle Biogenesis. J. Virol. 91, e00298–17 (2017).

20. Kakker, N. K., Mikhailov, M. V., Nermut, M. V., Burny, A. & Roy, P. Bovine leukemia virus Gag particle assembly in insect cells: formation of chimeric particles by domain-switched leukemia/lentivirus Gag polyprotein. Virology 265, 308–318 (1999).

21. Accola, M. A., Höglund, S. & Göttlinger, H. G. A putative alpha-helical structure which overlaps the capsid-p2 boundary in the human immunodeficiency virus type 1 Gag precursor is crucial for viral particle assembly. J. Virol. 72, 2072–2078 (1998).

22. Schur, F. K. M. et al. An atomic model of HIV-1 capsid-SP1 reveals structures regulating assembly and maturation. Science 353, 506–508 (2016).

23. Wagner, J. M. et al. Crystal structure of an HIV assembly and maturation switch. eLife 5, e17063 (2016).

24. Melamed, D. et al. The conserved carboxy terminus of the capsid domain of human immunodeficiency virus type 1 gag protein is important for virion assembly and release. J. Virol. 78, 9675–9688 (2004).

25. Dick, R. A. et al. Inositol phosphates are assembly co-factors for HIV-1. Nature 560, 509–512 (2018).

26. Briggs, J. a. G., et al. Structure and assembly of immature HIV. Proc. Natl. Acad. Sci. U. S. A. 106, 11090–11095 (2009).

27. de Marco, A. et al. Conserved and variable features of Gag structure and arrangement in immature retrovirus particles. J. Virol. 84, 11729–11736 (2010).

28. Bharat, T. A. M. et al. Cryo-electron microscopy of tubular arrays of HIV-1 Gag resolves structures essential for immature virus assembly. Proc. Natl. Acad. Sci. U. S. A. 111, 8233–8238 (2014).

29. Punjani, A. & Fleet, D. J. 3D variability analysis: Resolving continuous flexibility and discrete heterogeneity from single particle cryo-EM. J. Struct. Biol. 213, 107702 (2021).

30. Dodonova, S. O., Prinz, S., Bilanchone, V., Sandmeyer, S. & Briggs, J. A. G. Structure of the Ty3/Gypsy retrotransposon capsid and the evolution of retroviruses. Proc. Natl. Acad. Sci. U. S. A. 116, 10048–10057 (2019).

31. Calcraft, T. et al. Integrated cryoEM structure of a spumaretrovirus reveals cross-kingdom evolutionary relationships and the molecular basis for assembly and virus entry. Cell 187, 4213–4230.e19 (2024).

32. Klumpe, S. et al. In-cell structure and snapshots of copia retrotransposons in intact tissue by cryo-ET. Cell 188, 2094–2110.e18 (2025).

33. Renner, N. et al. HIV-1 is dependent on its immature lattice to recruit IP6 for mature capsid assembly. Nat. Struct. Mol. Biol. 30, 370–382 (2023).

34. Hadravová, R., et al. *In Vitro* Assembly of Virus-Like Particles of a Gammaretrovirus, the Murine Leukemia Virus XMRV. J. Virol. 86, 1297–1306 (2012).

35. Borsetti, A., Öhagen, Å. & Göttlinger, H. G. The C-Terminal Half of the Human Immunodeficiency Virus Type 1 Gag Precursor Is Sufficient for Efficient Particle Assembly. J. Virol. 72, 9313–9317 (1998).

36. Temple, J., Tripler, T. N., Shen, Q. & Xiong, Y. A snapshot of HIV-1 capsid-host interactions. Curr. Res. Struct. Biol. 2, 222–228 (2020).

37. Bester, S. M. et al. Structural and mechanistic bases for a potent HIV-1 capsid inhibitor. Science 370, 360–364 (2020).

38. Meuser, M. E. et al. A Hybrid Gag Lattice as a Structural Intermediate in HIV-1 Maturation. 2026.01.10.698794 Preprint at 10.64898/2026.01.10.698794 (2026).

39. Price, A. J. et al. Host cofactors and pharmacologic ligands share an essential interface in HIV-1 capsid that is lost upon disassembly. PLoS Pathog. 10, e1004459 (2014).

40. Yang, H. et al. Structural insight into HIV-1 capsid recognition by rhesus TRIM5α. Proc. Natl. Acad. Sci. U. S. A. 109, 18372–18377 (2012).

41. Gamble, T. R. et al. Crystal structure of human cyclophilin A bound to the amino-terminal domain of HIV-1 capsid. Cell 87, 1285–1294 (1996).

42. Punjani, A., Rubinstein, J. L., Fleet, D. J. & Brubaker, M. A. cryoSPARC: algorithms for rapid unsupervised cryo-EM structure determination. Nat. Methods 14, 290–296 (2017).

43. Punjani, A., Zhang, H. & Fleet, D. J. Non-uniform refinement: adaptive regularization improves single-particle cryo-EM reconstruction. Nat. Methods 17, 1214–1221 (2020).

44. Zivanov, J., Nakane, T. & Scheres, S. H. W. A Bayesian approach to beam-induced motion correction in cryo-EM single-particle analysis. IUCrJ 6, 5–17 (2019).

45. Croll, T. I. ISOLDE: a physically realistic environment for model building into low-resolution electron-density maps. Acta Crystallogr. Sect. Struct. Biol. 74, 519–530 (2018).

46. Meng, E. C. et al. UCSF ChimeraX: Tools for structure building and analysis. Protein Sci. Publ. Protein Soc. 32, e4792 (2023).

47. Liebschner, D. et al. Macromolecular structure determination using X-rays, neutrons and electrons: recent developments in Phenix. Acta Crystallogr. Sect. Struct. Biol. 75, 861– 877 (2019).

