## Supplemental Tables for "Retroviral CA subdomains encode their immature assembly properties independently of strict subdomain co-pairing"

### SUPPLEMENTARY TABLES

Table S1: *In vitro* assembly conditions

| ***Construct*** | ***In vitro assembly buffer*** | ***Incubation*** | ***Protein concentration*** |
|---|---|---|---|
| HTLV-1 CA (immature) | 50 mM MES, 200 mM NaCl, pH 6 | 4 ˚C for 4h, 26 ˚C O/N | 680 µM |
| HIV-1 CA-SP1 (immature) | 50 mM MES, 500 µM IP6, pH 6 | Dialysis at 4 ˚C for 4h | 500 µM |
| HIV-1 CA (mature) | 50 mM MES, 40 mM NaCl, 150 µM IP6, 5 mM ßme, pH 6 | 37 ˚C for 1.5h | 150 µM |

**Table S2: CryoEM data collection, processing, and refinement table**

| | CH1 (EMD-59880) (PDB 34IT) | CH1dl (EMD-59881) (PDB 34IU) | CH1-NTDm (EMD-59882) |
|---|---|---|---|
| **Data collection and processing** | | | |
| Microscope | Glacios cryo-TEM | Titan Krios G3i cryo-TEM | Glacios cryo-TEM |
| Magnification | 92,000 | 81,000 | 120,000 |
| Voltage (kV) | 200 | 300 | 200 |
| Electron exposure (e-/Å^2) | 50 | 80 | 50 |
| Defocus range (μm) | -1.2 to -1.8 | -1.2 to -1.8 | -1.2 to -1.8 |
| Pixel size (Å) | 1.511 | 1.089 | 1.182 |
| Symmetry imposed | C6 | C6 | C6 |
| Micrographs for SPA | 2,847 | 5,331 | 1,563 |
| Initial extracted particle images | 4,637,708 | 10,546,456 | 2,347,026 |
| Final particle images | 104,870 | 64,156 | 21,922 |
| Map resolution (Å) | 4.07 | 3.75 | 7.23 |
| FSC threshold | 0.143 | 0.143 | 0.143 |
| Map resolution range (Å) | 2.46 to 5.28 | 2.41 to 5.47 | 5.07 to 14.52 |
| **Refinement** | | | |
| Initial model used (PDB code) | 9N92/5L93 | 9N92/5L93 | - |
| Model resolution (Å) | 6.2 | 4 | - |
| FSC threshold | 0.5 | 0.5 | - |
| Map sharpening B factor (Å^2) | -143.5 | -101.8 | - |
| *Model composition* | | | |
| Non-hydrogen atoms | 29,394 | 30,096 | - |
| Protein residues | 3,780 | 3,870 | - |
| *B factors (Å^2)* | | | |
| Protein | 93.14/205.43/136.43 | 41.07/129.76/71.01 | - |
| *R.m.s. deviations* | | | |
| Bond lengths (Å) | 0.003 | 0.003 | - |
| Bond angles (˚) | 0.733 | 0.690 | - |
| *Validation* | | | |
| MolProbity score | 1.67 | 1.33 | - |
| Clashscore | 11.82 | 6.01 | - |
| Poor rotamers (%) | 0 | 0 | - |
| *Ramachandran plot* | | | |
| Favored (%) | 97.60 | 99.06 | - |
| Allowed (%) | 2.40 | 0.94 | - |
| Disallowed (%) | 0 | 0 | - |

**Table S3: RMSD values of 3DVA extremes relative to WT (PDB 9N92 for CA-NTD and 5L93 for CA-CTD) for hexamer and 18-mer clusters**

| | RMSD HTLV-1 hexamer (NTD) | RMSD HTLV-1 18-mer (NTD) | RMSD HIV-1 hexamer (CTD-SP1) | RMSD HIV-1 18-mer (CTD-SP1) |
|---|---|---|---|---|
| 3DVA comp. 1.1 | 2.0 | 3.0 | 1.1 | 1.7 |
| 3DVA comp. 1:2 | 2.0 | 3.2 | 1.1 | 1.7 |
| 3DVA comp. 2:1 | 2.3 | 3.5 | 1.1 | 2.1 |
| 3DVA comp. 2:2 | 2.2 | 3.3 | 1.1 | 1.9 |
| 3DVA comp. 3:1 | 2.2 | 3.4 | 1.1 | 2.0 |
| 3DVA comp. 3:2 | 2.2 | 3.3 | 1.1 | 1.9 |
| **Mean** | **2.1** | **3.3** | **1.1** | **1.9** |

**Table S4:** Displacement of helices 4 in CH1 and the 3DVA extremes relative to WT for the trimer interface measured by aligning one helix 4 of the trimer and calculating the average distance of each residue to wt for the remaining helices 4 (e.g., alignment of H4:A1, measurement on H4:A2 and H4:A3, where A H4 denotes helix 4 and A1, A2, A3 denote the respective helices of the trimer interface at interface A). The table displays the average distance for each measured helix 4. The intra-interface asymmetry is the absolute value of the displacement of the H4:2 and H4:3 for all interfaces (green <0.5 Å, yellow <1.0 Å, red >1.0 Å).

| | Helix measured | CH1 | CH1dl | 3DVA comp. 1.1 | 3DVA comp. 1:2 | 3DVA comp. 2:1 | 3DVA comp. 2:2 | 3DVA comp. 3:1 | 3DVA comp. 3:2 |
|---|---|---|---|---|---|---|---|---|---|
| Interface A | H4:A2 | 3.3 | 3.9 | 3.1 | 3.5 | 3.8 | 3.7 | 2.6 | 3.7 |
| | H4:A3 | 2.9 | 3.7 | 3.6 | 3.2 | 3.7 | 3.4 | 4.5 | 2.9 |
| **Calculated metrics** | **Interface Mean** | **3.1** | **3.8** | **3.3** | **3.3** | **3.8** | **3.6** | **3.6** | **3.3** |
| | **Intra-interface asymmetry** | - | - | **0.6** | **0.3** | **0.1** | **0.4** | **1.9** | **0.8** |
| Interface B | H4:B2 | 3.1 | 3.6 | 3.4 | 3.3 | 4.3 | 2.7 | 3.8 | 3.3 |
| | H4:B3 | 3.1 | 3.7 | 3.4 | 3.5 | 2.2 | 4.1 | 2.9 | 3.9 |
| **Calculated metrics** | **Interface Mean** | **3.1** | **3.7** | **3.4** | **3.4** | **3.2** | **3.4** | **3.3** | **3.6** |
| | **Intra-interface asymmetry** | - | - | **0.0** | **0.3** | **2.2** | **1.4** | **0.9** | **0.6** |
| Interface C | H4:C2 | 3.1 | 3.7 | 3.1 | 2.8 | 2.6 | 3.5 | 4.1 | 2.9 |
| | H4:C3 | 2.8 | 3.5 | 3.3 | 3.6 | 4.6 | 2.7 | 2.5 | 3.7 |
| **Calculated metrics** | **Interface Mean** | **3.0** | **3.6** | **3.2** | **3.2** | **3.6** | **3.1** | **3.3** | **3.3** |
| | **Intra-interface asymmetry** | - | - | **0.3** | **0.7** | **2.0** | **0.8** | **1.6** | **0.8** |
| Interface D | H4:D2 | 3.1 | 3.7 | 3.2 | 3.5 | 3.7 | 3.9 | 2.5 | 3.7 |
| | H4:D3 | 3.0 | 3.7 | 3.4 | 2.9 | 3.5 | 3.2 | 4.4 | 2.7 |
| **Calculated metrics** | **Interface Mean** | **3.0** | **3.7** | **3.3** | **3.2** | **3.6** | **3.5** | **3.5** | **3.2** |
| | **Intra-interface asymmetry** | - | - | **0.3** | **0.6** | **0.3** | **0.7** | **1.9** | **1.0** |
| Interface E | H4:E2 | 3.0 | 3.6 | 3.1 | 2.9 | 3.9 | 2.6 | 3.8 | 3.0 |
| | H4:E3 | 2.8 | 3.6 | 3.2 | 3.3 | 2.2 | 3.8 | 3.1 | 3.6 |
| **Calculated metrics** | **Interface Mean** | **2.9** | **3.6** | **3.1** | **3.1** | **3.0** | **3.2** | **3.4** | **3.3** |
| | **Intra-interface asymmetry** | - | - | **0.1** | **0.4** | **1.7** | **1.2** | **0.7** | **0.6** |
| Interface F | H4:F2 | 2.9 | 3.5 | 3.1 | 2.7 | 2.7 | 3.6 | 4.1 | 2.8 |
| | H4:F3 | 2.8 | 3.5 | 3.1 | 3.2 | 4.4 | 2.7 | 2.3 | 3.5 |
| **Calculated metrics** | **Interface Mean** | **2.8** | **3.5** | **3.1** | **3.0** | **3.6** | **3.2** | **3.2** | **3.2** |
| | **Intra-interface asymmetry** | - | - | **0.0** | **0.5** | **1.7** | **0.9** | **1.8** | **0.7** |
| ***All interfaces*** | ***Mean RMSD*** | **3.0** | **3.7** | **3.2** | **3.2** | **3.5** | **3.3** | **3.4** | **3.3** |

**Table S5:** Displacement of helix 9 in CH1 and the 3DVA extremes relative to WT for the dimer interface, measured by aligning one helix 9 of the dimer and calculating the average distance of each residue to wt for the remaining helix 9 (e.g., alignment of H9:A1, measurement on H9:A2 where H9 denotes helix 9 and A1, A2 denote the respective helices of the trimer interface at interface A). The table displays the average distance for each measured helix 9.

| | Helix measured | CH1 | CH1dl | 3DVA comp. 1.1 | 3DVA comp. 1:2 | 3DVA comp. 2:1 | 3DVA comp. 2:2 | 3DVA comp. 3:1 | 3DVA comp. 3:2 |
|---|---|---|---|---|---|---|---|---|---|
| Interface A | H9:A2 | 0.9 | 1.4 | 0.9 | 0.9 | 1.0 | 0.9 | 1.2 | 0.8 |
| Interface B | H9:B2 | 0.9 | 1.4 | 0.9 | 1.0 | 0.7 | 1.1 | 1.0 | 0.9 |
| Interface C | H9:C2 | 0.9 | 1.4 | 0.9 | 1.0 | 1.1 | 0.9 | 0.8 | 1.0 |
| Interface D | H9:D2 | 0.9 | 1.4 | 0.9 | 0.9 | 1.0 | 0.9 | 1.2 | 0.8 |
| Interface E | H9:E2 | 0.9 | 1.4 | 0.9 | 0.9 | 0.7 | 1.1 | 1.0 | 0.9 |
| Interface F | H9:F2 | 0.9 | 1.4 | 0.9 | 0.9 | 1.2 | 0.9 | 0.8 | 1.0 |
| ***All interfaces*** | ***Mean RMSD*** | ***0.9*** | ***1.4*** | ***0.9*** | ***0.9*** | ***1.0*** | ***1.0*** | ***1.0*** | ***0.9*** |
